# Identifying disease-critical cell types and cellular processes across the human body by integration of single-cell profiles and human genetics

**DOI:** 10.1101/2021.03.19.436212

**Authors:** Karthik A. Jagadeesh, Kushal K. Dey, Daniel T. Montoro, Rahul Mohan, Steven Gazal, Jesse M. Engreitz, Ramnik J. Xavier, Alkes L. Price, Aviv Regev

## Abstract

Genome-wide association studies (GWAS) provide a powerful means to identify loci and genes contributing to disease, but in many cases the related cell types/states through which genes confer disease risk remain unknown. Deciphering such relationships is important for identifying pathogenic processes and developing therapeutics. Here, we introduce sc-linker, a framework for integrating single-cell RNA-seq (scRNA-seq), epigenomic maps and GWAS summary statistics to infer the underlying cell types and processes by which genetic variants influence disease. We analyzed 1.6 million scRNA-seq profiles from 209 individuals spanning 11 tissue types and 6 disease conditions, and constructed gene programs capturing cell types, disease progression, and cellular processes both within and across cell types. We evaluated these gene programs for disease enrichment by transforming them to SNP annotations with tissue-specific epigenomic maps and computing enrichment scores across 60 diseases and complex traits (average *N=*297K). Cell type, disease progression, and cellular process programs captured distinct heritability signals even within the same cell type, as we show in multiple complex diseases that affect the brain (Alzheimer’s disease, multiple sclerosis), colon (ulcerative colitis) and lung (asthma, idiopathic pulmonary fibrosis, severe COVID-19). The inferred disease enrichments recapitulated known biology and highlighted novel cell-disease relationships, including GABAergic neurons in major depressive disorder (MDD), a disease progression M cell program in ulcerative colitis, and a disease-specific complement cascade process in multiple sclerosis. In autoimmune disease, both healthy and disease progression immune cell type programs were associated, whereas for epithelial cells, disease progression programs were most prominent, perhaps suggesting a role in disease progression over initiation. Our framework provides a powerful approach for identifying the cell types and cellular processes by which genetic variants influence disease.

## INTRODUCTION

Genome wide association studies (GWAS) have successfully identified thousands of disease-associated variants^1–3^, but the cellular mechanisms through which these variants drive complex diseases and traits remain largely unknown. This is due to several challenges, including the difficulty of relating the approximately 95% of risk variants that reside in non-coding regulatory regions to the genes they regulate^4–7^, and our limited knowledge of the specific cells and functional programs in which these genes are active^8^. Previous studies have linked traits to functional elements^9–15^ and to cell types from bulk RNA-seq profiles^16–18^. Considerable work remains to analyze cell types and states at finer resolutions across a breadth of tissues, incorporate disease tissue-specific gene expression patterns, model cellular processes within and across cell types, and leverage enhancer-gene links^19–23^ to improve power.

ScRNA-seq data provide a unique opportunity to tackle these challenges^24^. Single-cell profiles allow the construction of multiple gene programs to more finely relate GWAS variants to function, including programs that reflect cell-type-specific signatures^25–28^, disease progression within cell types^29, 30^, and key cellular processes that vary within and/or across cell types^31^. Initial studies have related single-cell profiles with human genetics in *post hoc* analyses by mapping candidate genes from disease-associated genomic regions to cell types by their expression relative to other cell types^32–34^. More recent studies have begun to leverage genome-wide polygenic signals to map traits to cell types from single cells within the context of a single tissue^35–37^. However, focusing on a single tissue could in principle result in misleading conclusions, because disease mechanisms span tissue types across the human body. For example, in the context of the colon, a neural gene associated with psychiatric disorders would appear highly specific to enteric neurons, but this cell population may no longer be strongly implicated when the analysis also includes cells from the human central nervous system (CNS)^38^. Thus, there is a need for a principled method that combines human genetics and comprehensive scRNA-seq applied across multiple tissues and organs.

Here, we develop and apply sc-linker, an integrated framework to relate human disease and complex traits to cell types and cellular processes by integrating GWAS summary statistics, epigenomics and scRNA-seq data from multiple tissue types, diseases, individuals and cells. Unlike previous studies, we analyze gene programs that represent different facets of cells, including discrete types, processes activated specifically in a cell type in disease, and gene programs that vary across cells irrespective of cell type definitions (recovered by latent factor models). We transform gene programs to SNP annotations using tissue-specific enhancer-gene links^19–23^ in preference to standard gene window-based linking strategies used in existing gene-set enrichment methods such as MAGMA^39^, RSS-E^13^ and LDSC-SEG^18^. We then link SNP annotations to diseases by applying stratified LD score regression^11^ (S-LDSC) with the baseline-LD model^40, 41^ to the resulting SNP annotations. We further integrate cellular expression and GWAS to prioritize specific genes in the context of disease-critical gene programs, thus shedding light on underlying disease mechanisms.

## RESULTS

### Overview of sc-linker

We developed a framework to link gene programs derived from scRNA-seq with diseases and complex traits (**Fig. 1a**). First, we use scRNA-seq to construct gene programs, defined as probabilistic gene sets, that characterize (1) individual cell types, (2) disease progression (disease *vs.* healthy cells of the same type), or (3) cellular processes (**Methods**). Then, we link the genes underlying these programs to SNPs that regulate them by incorporating two tissue-specific enhancer-gene linking strategies: Roadmap Enhancer-Gene Linking^19–21^ and the Activity-by-Contact (ABC) model^22, 23^. Finally, we evaluate the disease informativeness of the resulting SNP annotations by applying S-LDSC^11^ conditional on a broad set of coding, conserved, regulatory and LD-related annotations from the baseline-LD model^40, 41^. Altogether, our approach links diseases and traits with gene programs recapitulating cell types and cellular processes. We have released open-source software implementing the approach (sc-linker; **Code Availability**), a web interface for visualizing the results (**Data Availability**), postprocessed scRNA-seq data, gene programs, enhancer-gene linking strategies, and SNP annotations analyzed in this study (**Data Availability**).

**Fig. 1.**
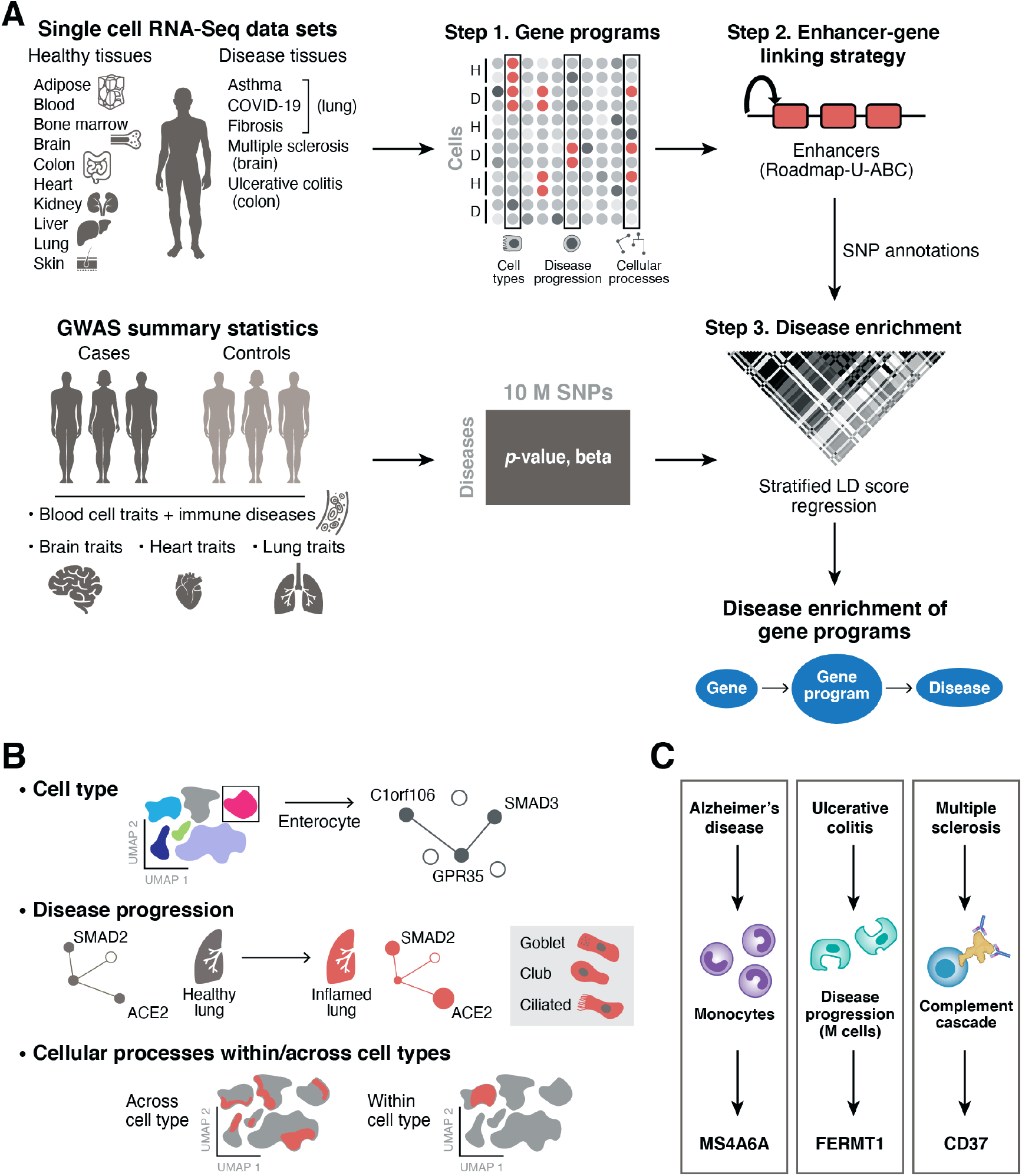
Approach for identifying disease-critical cell types and cellular processes by integration of single-cell profiles and human genetics. **a**. sc-linker framework. Left: Input. scRNA-seq (top) and GWAS (bottom) data. Middle and right: Step 1: Deriving cell type, disease progression, and cellular process gene programs from scRNA-seq (top) and associating SNPs with traits from human GWAS (bottom). Step 2: Generation of SNP annotations. Gene programs are linked to SNPs by enhancer-gene linking strategies to generate SNP annotations. Step 3: S-LDSC is applied to the resulting SNP annotations to evaluate heritability enrichment for a trait. **b.** Constructing gene programs. Top: Cell type programs of genes specifically expressed in one cell type *vs*. others. Middle: disease progression programs of genes specifically expressed in cells of the same type in disease *vs*. healthy samples. Bottom: cellular process programs of genes co-varying either within or across cell subsets; these programs may be healthy-specific, disease-specific, or shared. **c.** Examples of disease-gene program-gene relationships recovered by our framework.

We constructed three kinds of gene programs from scRNA-seq data (**Fig. 1b**): (i) cell type programs that represent genes specifically enriched in an individual broad cell type of a tissue (*e.g.*, colon T cells) compared to other cell types in that tissue; (ii) disease progression cell type programs that represent disease progression differences in gene expression within the same cell type (*e.g.*, colon T cells in UC *vs*. healthy colon); and (iii) cellular process programs that capture gene co-variation patterns within and across cell types (*e.g.*, MHC class II antigen presenting process varying across dendritic cells and B cells) (**Methods**). We constructed (healthy) cell type programs by assessing the differential expression of each gene for the focal cell type *vs*. all other cell types in the tissue using healthy individuals (with cell types defined by clustering^42^ and annotated *post hoc*) and transforming each gene’s Z score to a probabilistic score (**Methods**). (Analogous to healthy cell type programs generated from healthy tissues, we also generated disease cell type programs from cell profiles from disease tissues.) We constructed disease progression cell type programs by assessing differential expression between cells of the same type in disease *vs*. healthy tissue and transforming each gene’s Z score to a probabilistic score (**Methods**), aiming to capture genes involved in disease progression and symptoms after onset. (We caution that disease progression programs may also capture genes reflecting genetic susceptibility to disease, rather than progression.) On average, disease progression cell type programs had low correlation with healthy cell type programs of the same cell type (Pearson *r*=0.16 across tissues; see below) compared to the much higher correlation between disease and healthy cell type programs (average *r*=0.62 across tissues); thus, we did not consider disease cell type programs in any of our primary analyses. Finally, independently of predefined cell type subsets, we constructed cellular process programs using unsupervised learning, via non-negative matrix factorization^43^ (NMF) and a modified NMF (to jointly model both healthy and disease states) of normalized gene expression values, with the latent factors (programs) representing variation across continuums of cell types or processes active in multiple cell types. We computed the correlations between weights of each latent factor across cells and each gene’s expression across cells and then transformed them to a 0-1 probabilistic scale to define each cellular process program. We annotated each cellular process program by its most enriched pathways (**Methods**) and labeled it as ‘intra-cell type’ or ‘inter-cell type’ if highly correlated with only one or multiple cell type programs, respectively (**Methods**). Intra-cell type cellular processes can correspond to narrower cell types (*e.g.*, CD4 T cells) reflecting cell subsets of broader cell type categories (*e.g.*, T cells) or variation within a cell type continuum, whereas inter-cell type cellular process programs can reflect shared processes or transitions.

Next, we transformed the genes prioritized by each program into SNP annotations by linking each gene to SNPs that may regulate their activity in *cis* (**Fig. 1a**). We generated SNP annotations using an enhancer-gene linking strategy, defined as an assignment of 0, 1 or more linked genes to each SNP, combining Roadmap Enhancer-Gene Linking (Roadmap)^19, 21^ and Activity-By-Contact (ABC)^22, 23^ strategies (Roadmap∪ABC) in the tissue underlying the program of interest (**Methods**). We used tissue level enhancer-gene links instead of cell type level enhancer-gene links because they generated more significant associations in benchmarking experiments based on current data (see below). We primarily focused on linking genes to non-coding regulatory variants (which may drive cell-type specific differences in expression), based on the results of our benchmarking experiments (see below).

Finally, we evaluated each gene program for disease heritability enrichment by applying S-LDSC^11^ with the baseline-LD model^40, 41^ to the resulting SNP annotations (**Fig. 1a**, **Methods**). The S-LDSC analysis was conditioned on 86 coding, conserved, regulatory and LD-related annotations from the baseline-LD model (v2.1)^40, 41^ (**Data Availability**), and uses heritability enrichment to evaluate informativeness for disease. Heritability enrichment is defined as the proportion of heritability explained by SNPs in an annotation divided by the proportion of SNPs in the annotation^11^; this generalizes to annotations with values between 0 and 1^44^. We further define the Enrichment score (E-score) of a gene program as the difference between the heritability enrichment of the SNP annotation corresponding to the gene program of interest and the SNP annotation corresponding to a gene program assigning a probabilistic grade of 1 to all protein-coding genes with at least one enhancer-gene link in the relevant tissue (**Methods**). We use the p-value of the E-score as our primary metric, assessing statistical significance using a genomic block-jackknife as in our previous work^11^, because the p-values can be compared across datasets, whereas the E-score magnitude can vary substantially in gene programs dominated by a smaller (or larger) number of genes. We primarily focus on E-scores greater than 2, because E-scores that are statistically significant but small in magnitude may have more limited biological importance, as the cell types underlying these E-scores may be tagging other causal cell types. We performed this analysis over healthy cell type programs (**data file S1**), disease progression programs (**data file S2**), and cellular process programs (**data file S3**). We identified the top 50 genes driving disease enrichments with highest proximity based MAGMA (v 1.08) gene-disease association scores^39^ of genes with high probabilistic grade in each gene program (**Fig. 1C, data file S4, Methods**) focusing on genes that are both (i) close to a GWAS signal and (ii) in an enriched gene program.

We analyzed a broad range of human scRNA-seq data, spanning 17 data sets from 11 tissues and 6 disease conditions. The 11 non-disease tissues include immune (peripheral blood mononuclear cells (PBMCs)^26, 45^, cord blood^27^, and bone marrow^27^), brain^28^, kidney^46^, liver^47^, heart^25^, lung^29^, colon^34^, skin^48^ and adipose^47^. The 6 disease conditions include multiple sclerosis (MS) brain^49^, Alzheimer’s disease brain^30^, ulcerative colitis (UC) colon^34^, asthma lung^50^, idiopathic pulmonary fibrosis (IPF) lung^29^ and COVID-19 bronchoalveolar lavage fluid^51^ (**Supplementary Fig. 1**). In total, the scRNA-seq data includes 209 individuals, 1,602,614 cells and 256 annotated cell subsets (**Methods**, **Supplementary Table 1**). We also compiled publicly available GWAS summary statistics for 60 unique diseases and complex traits (genetic correlation < 0.9; average *N*=297K) (**Methods**, **Supplementary Table 2**). We analyzed gene programs from each scRNA-seq dataset in conjunction with each of 60 diseases and complex traits, but we primarily report those that are most pertinent for each program.

### Benchmarking sc-linker

As a proof of principle, we benchmarked sc-linker by analyzing 5 blood cell traits that inherently correspond to underling cell types (**Supplementary Table 2**) using immune cell type programs constructed from scRNA-seq data (**Fig. 2a,b**, **Supplementary Fig. 1**). We constructed 6 immune cell type programs that were identified across 4 data sets – two from PBMCs (*k=*4,640 cells; *n*=2 individuals^26^; *k=*68,551; *n*=8 individuals^45^), and one each of cord blood^27^ (*k=*263,828; *n*=8) and bone marrow^27^ (*k=*283,894; *n*=8). We identified the expected cell type enrichments, including enrichment of erythroid cells for red blood cell count, megakaryocytes for platelet count, monocytes for monocyte count, and of B cells and T cells for lymphocyte percentage (**Fig. 2d**, **Supplementary Fig. 2a**).

**Fig. 2.**
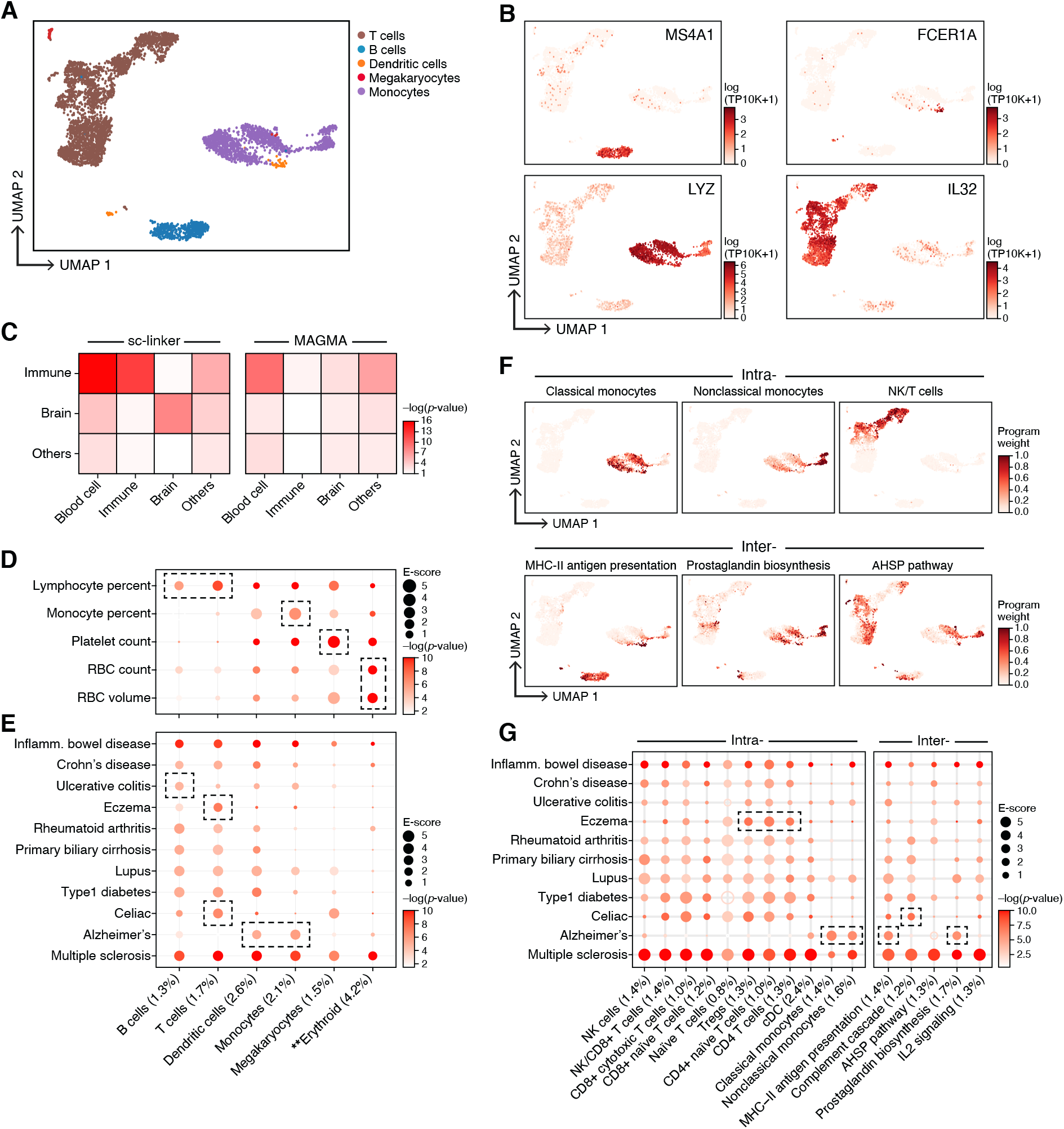
Linking immune cell types and cellular processes to immune-related diseases and blood cell traits. **a,b**. Immune cell types. Uniform Manifold Approximation and Projection (UMAP) embedding of peripheral blood mononuclear cell (PBMC) scRNA-seq profiles (dots) colored by cell type annotations (a) or expression of cell-type-specific genes (b). **c.** Benchmarking of sc-linker *vs*. MAGMA. Significance (average -log_10_(p-value)) of association between immune, brain and other tissue cell type programs (rows) and blood cell, immune-related, brain-related and other traits (columns) for sc-linker (left) and MAGMA gene set analysis (right). Other cell types x other diseases/traits are not included in the specificity calculation, due to the broad set of cell types and diseases/traits in this category. **d,e.** Enrichments of immune cell type programs for blood cell traits and immune-related diseases. Magnitude (E-score, dot size) and significance (-log_10_(P-value), dot color) of the heritability enrichment of immune cell type programs (columns) for blood cell traits (rows, d) or immune-related diseases (rows, e). **f.** Examples of inter- and intra-cell type cellular process programs. UMAP of PBMC (as in a), colored by each program weight (color bar) from non-negative matrix factorization (NMF). **g.** Enrichments of immune cellular process programs for immune-related diseases. Magnitude (E-score, dot size) and significance (-log_10_(p-value), dot color) of the heritability enrichment of cellular process programs (columns) for immune-related diseases (rows). In panels d,e,g, the size of each corresponding SNP annotation (% of SNPs) is reported in parentheses. Numerical results are reported in **data file S1,3**. Further details of all diseases and traits analyzed are provided in **Supplementary Table 2**. **Erythroid cells were observed in only bone marrow and cord blood datasets.

The Roadmap∪ABC enhancer-gene linking strategy outperformed every other enhancer-gene linking strategy we tested in identifying these expected enrichments, including its constituent Roadmap and ABC strategies, the standard 100kb window-based approach used in LDSC-SEG^18^ (**Supplementary Fig. 3d,e**), other gene-proximal SNP-gene linking strategies such as exons and closest TSS at different genomic distances, other regulatory SNP-gene linking strategies using Promoter-capture Hi-C and eQTLs in GTEx whole blood, and combinations of these strategies (**Supplementary Fig. 3a,b and data file S5**). Additionally, the tissue-specific Roadmap∪ABC-immune enhancer-gene linking strategy generated slightly higher specificity of cell type enrichments compared to cell-type-specific enhancer-gene linking strategies, supporting the use of tissue-specific enhancer-gene linking (**Supplementary Fig. 4l**). This trend may stem from existing cell-type-specific enhancer-gene links being noisier, due to the limited amount of underlying cell-type-specific data, or because tissue-specific enhancer-gene links may tag enhancer-gene links in causal cell types that were not assayed (distinct from tagging captured by cell type programs).

The cell type programs were robust to the number of cells and individuals. Specifically, cell type programs and their corresponding enrichment results were robust (correlation of *r*=0.91) to changes in the number of profiled cells for scRNA-seq datasets with greater than 500 cells (**Supplementary Fig. 4f-h**); larger scRNA-seq datasets can uncover cell populations and states that may be missed in smaller datasets, due to sampling power. The cell type programs were also highly similar across different sets of individuals (*r*=0.96 on average between programs of the same cell type generated from different samples, with consistent specificity in expected enrichments; **Supplementary Fig. 4i-k**).

We observed higher specificity of enrichments of relevant cell type-trait pairs for our polygenic approach based on specifically expressed genes *vs*. other cell types compared to several other approaches including (i) functional enrichment of fine-mapped SNPs^52^ (**Supplementary Fig. 5a**); (ii) all expressed genes in a cell type, defined across several thresholds (**Supplementary Fig. 6**); (iii) specifically expressed genes *vs*. other genes in the same cell type; or (iv) specifically expressed genes *vs*. other genes in the same cell type, after normalizing each gene across cell types (**Supplementary Fig. 7 and data file S6**). We hypothesize that the “*all expressed genes*” approach greatly underperforms sc-linker because, for a given expressed gene, centrality of function in a cell is often reflected in its level of expression compared to that in other cells^53, 54^.

Sc-linker also outperformed three methods that use the MAGMA software^39^. First, we considered a baseline method for scoring cell types by scoring each cell using MAGMA *gene-level* associations to a trait and averaging across all cells of a cell type. We scored each cell for a trait using the top 200 MAGMA genes with highest score for the trait, computing the average expression over all the genes and subtracting an expression-matched control gene set. Sc-linker generated higher cell type specificity in enrichments compared to this baseline (**Supplementary Fig. 4n**). Second, sc-linker outperformed MAGMA *gene set-level* associations with a sensitivity/specificity index of 6.29 for sc-linker vs. 5.83 for MAGMA (**Supplementary Fig. 8**), and this was further underscored by a comparison across a broader set of cell types and diseases. Specifically, analyzing 3 major cell type categories (immune, brain, other) and 4 major categories of diseases (blood biomarkers, immune-related diseases, brain-related diseases, other diseases), and focusing on the most plausible pairings (immune cell types x blood biomarkers, immune cell types x immune-related diseases, and brain cell types x brain-related diseases), sc-linker attained a higher sensitivity/specificity index (9.47) compared to MAGMA (1.78) (**Methods, Fig. 2c**). This is consistent with prior work showing that MAGMA can produce significant results in the absence of true enrichment (false positives) due to uncorrected genomic confounding (*e.g.*, non-gene set-specific exon enrichment), if no gene-level covariates are included to correct for potential confounding. Third, FUMA^35^, a web interface that applies (gene set-level) MAGMA with precompiled scRNA-seq data (distinct from the data in our study), underperformed both sc-linker and gene set-level MAGMA (**data file S7 and S8**).

### Distinguishing innate, adaptive and antibody-mediated immunity contributions among immune-related diseases

We next analyzed 11 autoimmune and/or inflammation-associated diseases (**Supplementary Table 2**) using the 6 immune cell type programs above (**Fig. 2a,b**, **Supplementary Fig. 1**) and 10 (intra-cell type and inter-cell type) immune cellular process programs (**Fig. 2f**). (Enrichment results for the remaining 49 diseases and traits with immune cell type programs are reported in **Supplementary Fig. 9**; we did not construct disease progression programs, as these datasets included healthy samples only). We identified cell type-disease enrichments that conform to known disease biology (**Fig. 2e**, **Supplementary Fig. 2b**), including T cells for eczema^55, 56^, B and T cells for primary biliary cirrhosis (PBC)^18^, and dendritic cells and monocytes for Alzheimer’s disease^57^. Additionally, the highly significant enrichments for MS across all 6 immune cell type programs analyzed are consistent with previous analyses^18, 58–61^, supporting the validity of our approach.

Several of the significant cell type-disease enrichments are not as widely established and may implicate previously unexplored biological mechanisms (**Fig. 2e**, **Table 1, Supplementary Fig. 2b**). For example, we detected significant enrichment in B cells for UC; B cells have been detected in basal lymphoid aggregates in the ulcerative colitis (UC) colon, but their pathogenic significance remains unknown^62^. In addition, T cells were highly enriched for celiac disease; the top driving genes including *ETS1* (ranked 1), associated with T cell development and IL2 signaling^63^, and *CD28* (ranked 3), critical for T cell activation. This suggests that aberrant T cell maintenance and activation may impact inflammation in celiac disease. Recent reports of a permanent loss of resident gamma delta T cells in the celiac bowel and the subsequent recruitment of inflammatory T cells may further support this hypothesis^64^. These results were recapitulated across an independent immune cell scRNA-seq dataset, both in the gene programs (average correlation: 0.78 for the same cell type) and disease enrichments (0.86 correlation of the E-score over all cell type and trait pairs). A cross-trait analysis of the patterns of cell type enrichments suggests that Celiac disease and rheumatoid arthritis involves cell-mediated adaptive immune response, UC and primary biliary cirrhosis involve antibody-mediated adaptive immune response, Alzheimer’s disease has a strong signal of innate immune, and MS and IBD involve contributions from a wide range of immune cell types (**Supplementary Fig. 10**).

**Table 1.**
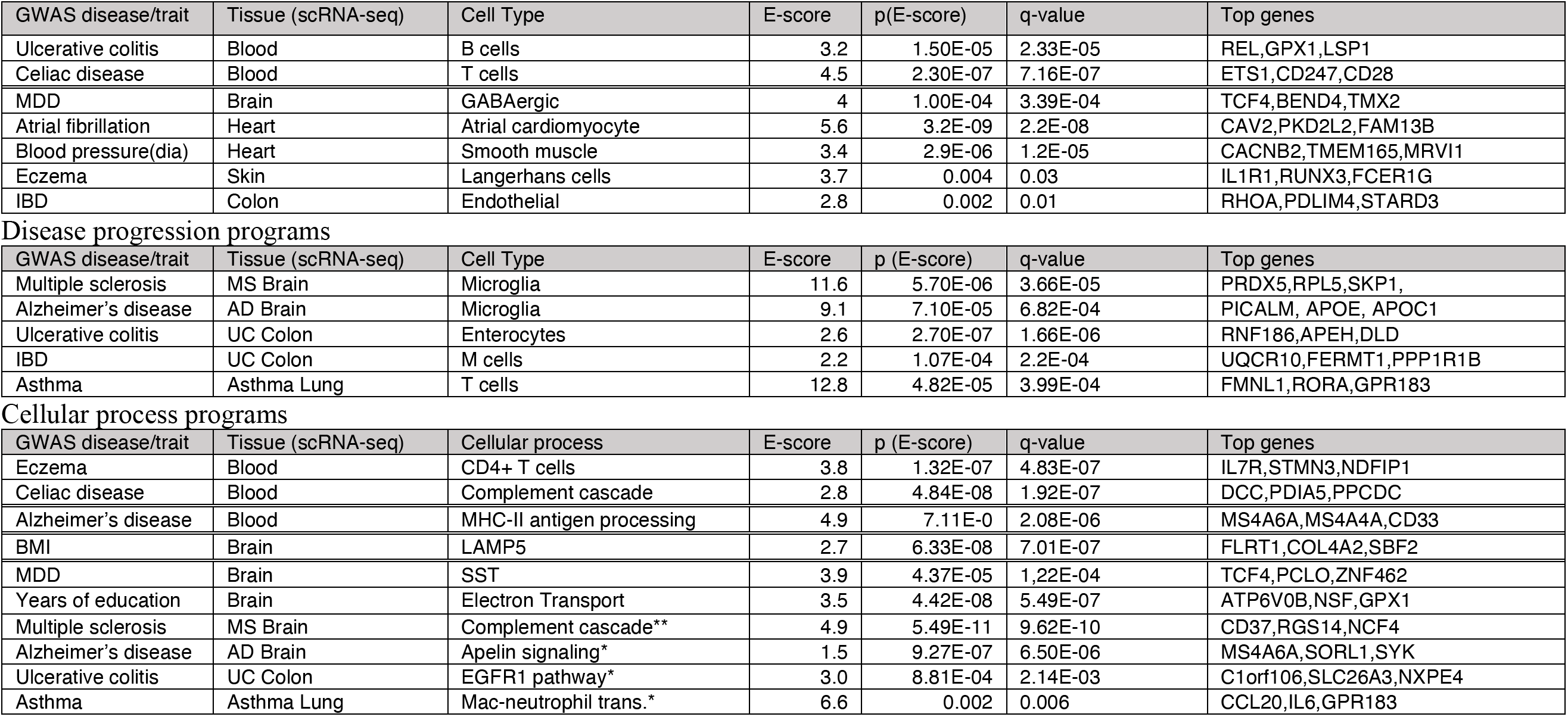
Notable enrichments from analyses of cell type, disease progression and cellular process gene programs. For each notable enrichment, we report the GWAS disease/trait, tissue source for scRNA-seq data, cell type, enrichment score (E-score), 1-sided p-value for positive E-score, and top genes driving the enrichment. Nominally significant enrichments for diseases with limited GWAS sample size are colored in grey. MDD is an abbreviation for major depressive disorder, blood pressure (dia.) is an abbreviation for diastolic blood pressure, mac-neutrophil trans. is an abbreviation for macrophage-neutrophil transition. * denotes cellular process programs shared across healthy and disease states. ** denotes cellular process programs specific to disease states. The full list of genes driving these associations is provided in **data file S4**.

Cell type programs
| GWAS disease/trait | Tissue (scRNA-seq) | Cell Type | E-score | p(E-score) | q-value | Top genes |
| --- | --- | --- | --- | --- | --- | --- |
| Ulcerative colitis | Blood | B cells | 3.2 | 1.50E-05 | 2.33E-05 | REL,GPX1,LSP1 |
| Celiac disease | Blood | T cells | 4.5 | 2.30E-07 | 7.16E-07 | ETS1,CD247,CD28 |
| MDD | Brain | GABAergic | 4 | 1.00E-04 | 3.39E-04 | TCF4,BEND4,TMX2 |
| Atrial fibrillation | Heart | Atrial cardiomyocyte | 5.6 | 3.2E-09 | 2.2E-08 | CAV2,PKD2L2,FAM13B |
| Blood pressure(dia) | Heart | Smooth muscle | 3.4 | 2.9E-06 | 1.2E-05 | CACNB2,TMEM165,MRVI1 |
| Eczema | Skin | Langerhans cells | 3.7 | 0.004 | 0.03 | IL1R1,RUNX3,FCER1G |
| IBD | Colon | Endothelial | 2.8 | 0.002 | 0.01 | RHOA,PDLIM4,STARD3 |

Disease progression programs
| GWAS disease/trait | Tissue (scRNA-seq) | Cell Type | E-score | p (E-score) | q-value | Top genes |
| --- | --- | --- | --- | --- | --- | --- |
| Multiple sclerosis | MS Brain | Microglia | 11.6 | 5.70E-06 | 3.66E-05 | PRDX5,RPL5,SKP1, |
| Alzheimer's disease | AD Brain | Microglia | 9.1 | 7.10E-05 | 6.82E-04 | PICALM, APOE, APOC1 |
| Ulcerative colitis | UC Colon | Enterocytes | 2.6 | 2.70E-07 | 1.66E-06 | RNF186,APEH,DLD |
| IBD | UC Colon | M cells | 2.2 | 1.07E-04 | 2.2E-04 | UQCR10,FERMT1,PPP1R1B |
| Asthma | Asthma Lung | T cells | 12.8 | 4.82E-05 | 3.99E-04 | FMNL1,RORA,GPR183 |

**Table 1. Notable enrichments from analyses of cell type, disease progression and cellular process gene programs.** For each notable
| GWAS disease/trait | Tissue (scRNA-seq) | Cellular process | E-score | p (E-score) | q-value | Top genes |
| --- | --- | --- | --- | --- | --- | --- |
| Eczema | Blood | CD4+ T cells | 3.8 | 1.32E-07 | 4.83E-07 | IL7R,STMN3,NDFIP1 |
| Celiac disease | Blood | Complement cascade | 2.8 | 4.84E-08 | 1.92E-07 | DCC,PDIA5,PPCDC |
| Alzheimer's disease | Blood | MHC-II antigen processing | 4.9 | 7.11E-0 | 2.08E-06 | MS4A6A,MS4A4A,CD33 |
| BMI | Brain | LAMP5 | 2.7 | 6.33E-08 | 7.01E-07 | FLRT1,COL4A2,SBF2 |
| MDD | Brain | SST | 3.9 | 4.37E-05 | 1.22E-04 | TCF4,PCLO,ZNF462 |
| Years of education | Brain | Electron Transport | 3.5 | 4.42E-08 | 5.49E-07 | ATP6V0B,NSF,GPX1 |
| Multiple sclerosis | MS Brain | Complement cascade** | 4.9 | 5.49E-11 | 9.62E-10 | CD37,RGS14,NCF4 |
| Alzheimer's disease | AD Brain | Apelin signaling* | 1.5 | 9.27E-07 | 6.50E-06 | MS4A6A,SORL1,SYK |
| Ulcerative colitis | UC Colon | EGFR1 pathway* | 3.0 | 8.81E-04 | 2.14E-03 | C1orf106,SLC26A3,NXPE4 |
| Asthma | Asthma Lung | Mac-neutrophil trans.* | 6.6 | 0.002 | 0.006 | CCL20,IL6,GPR183 |

Analyzing the 10 immune cellular process programs (**Fig. 2f**) across the 11 immune-related diseases and 5 blood cell traits, we identified both disease-specific enrichments and others shared across diseases (**Fig. 2g**, **Table 1**). For example, while T cells have been previously linked to eczema, we pinpointed higher enrichment in CD4^+^ T cells compared to CD8^+^ T cells. The IL2 signaling cellular process program in T and B cells was significantly enriched for both eczema and celiac disease, though the genes driving the enrichment were not significantly overlapping (p-value: 0.21). Additionally, the complement cascade cellular process program in plasma, B, and hematopoietic stem cells (HSCs) was most highly enriched among all inter cellular programs for celiac disease. For Alzheimer’s disease, there was a strong enrichment in both classical and non-classical monocyte intra-cell type cellular programs, and in MHC class II antigen presentation (inter cell type; dendritic cells (DCs) and B cells) and prostaglandin biosynthesis (inter cell type; monocytes, DCs, B cells and T cells) programs. Among the notable driver genes were: *IL7R* (ranked 1) and *NDFIP1* (ranked 3) for CD4^+^ T cells in eczema, which respectively play key roles in Th2 cell differentiation^65, 66^ and in mediating peripheral *CD4* T cell tolerance and allergic reactions^67, 68^; and *CD33* (ranked 1) in MHC class II antigen processing in Alzheimer’s disease, a microglial receptor strongly associated with increased risk in previous GWAS^69, 70^.

### Linking GABAergic and glutamatergic neurons to psychiatric disease

We next focused on brain cells and psychiatric disease, by analyzing 9 cell type programs (**Fig. 3a**) and 12 cell process programs (**Fig. 3e**, 10 intra- and 2 inter-cell type programs) from scRNA-seq data of brain prefrontal cortex (*k*=73,191, *n*=10)^28^ **(Supplementary Table 1**) with 11 psychiatric or neurological diseases and traits (**Supplementary Table 2**). We did not construct disease progression programs, as this dataset included healthy samples only.

**Fig. 3.**
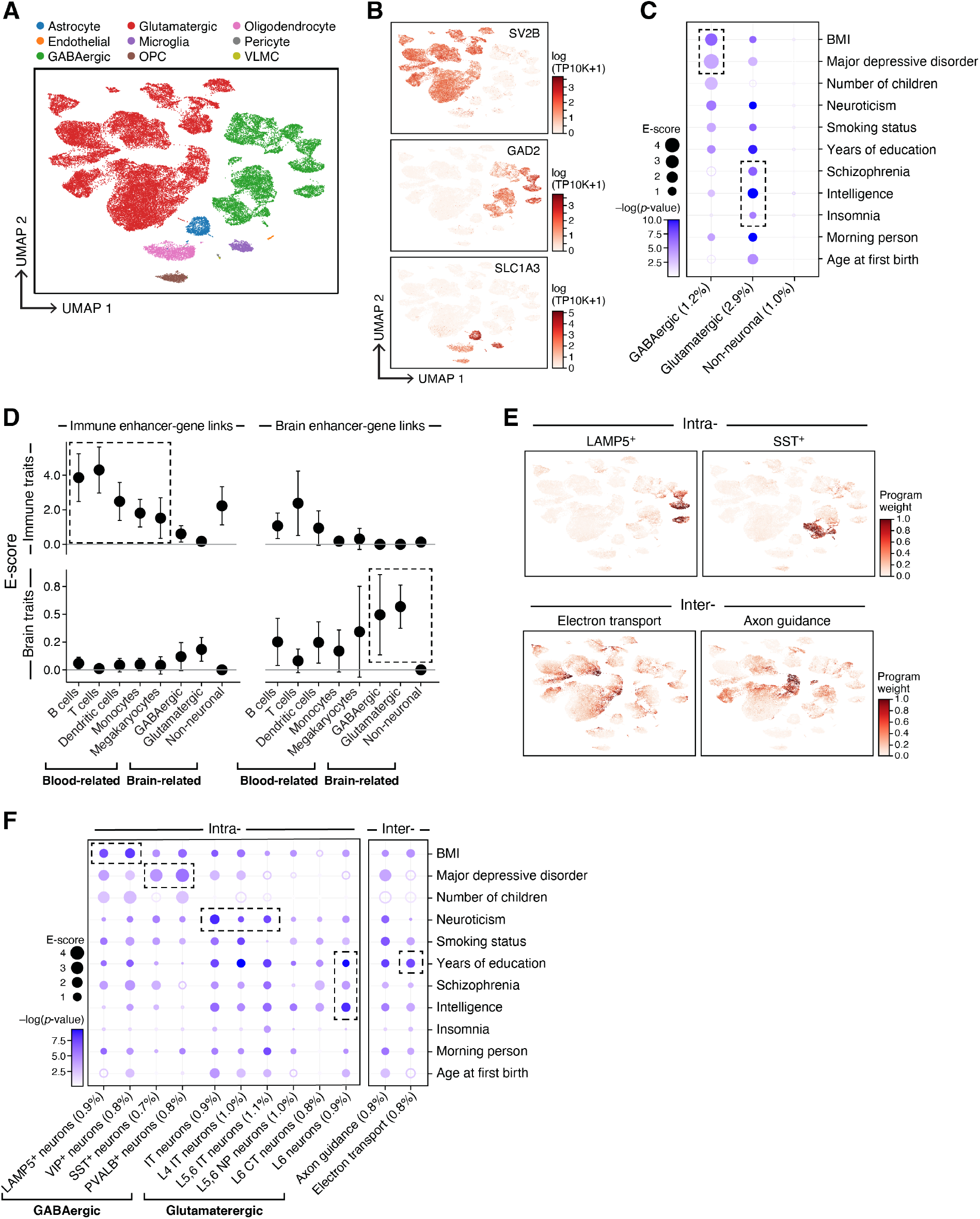
Linking neuron cell subsets and cellular processes to brain-related diseases and traits. **a,b.** Major brain cell types. UMAP embedding of brain scRNA-seq profiles (dots) colored by cell type annotations (a) or expression of cell-type-specific genes (b). **c.** Enrichments of brain cell type programs for brain-related diseases and traits. Magnitude (E-score, dot size) and significance (-log_10(_P-value), dot color) of the heritability enrichment of brain cell type programs (columns) for brain-related diseases and traits (rows). **d.** Comparison of immune vs. brain cell type programs, enhancer-gene linking strategies, and diseases/traits. Magnitude (E-score and SE) of the heritability enrichment of immune vs. brain cell type programs (columns) constructed using immune vs. brain enhancer-gene linking strategies (left and right panels) for immune-related vs. brain-related diseases and traits (top and bottom panels). **e.** Examples of inter- and intra-cell type cellular processes. UMAP (as in a), colored by each program weight (color bar) from non-negative matrix factorization (NMF). **f.** Enrichments of brain cellular process programs for brain-related diseases and traits. Each of the cellular process programs is constructed using NMF to decompose the cells by genes matrix into two matrices, cells by programs and programs by genes. Magnitude (E-score, dot size) and significance (-log_10_(P-value), dot color) of the heritability enrichment of cellular process programs (columns) for brain-related diseases and traits (rows). In panels c and f, the size of each corresponding SNP annotation (% of SNPs) is reported in parentheses. Numerical results are reported in **data file S1,3**. Further details of all diseases and traits analyzed are provided in **Supplementary Table 2**.

Notably, we observed enrichments of major depressive disorder (MDD) and body mass index (BMI) specifically in GABAergic neurons, while insomnia, schizophrenia (SCZ), and intelligence were highly enriched specifically in glutamatergic neurons, and neuroticism was highly enriched in both. GABAergic neurons regulate the brain’s ability to control stress levels, which is the most prominent vulnerability factor in MDD^71^ (**Fig. 3b,c**, **Table 1, Supplementary Fig. 2c**). Among the top genes driving this enrichment were *TCF4* (ranked 1), a critical component for neuronal differentiation that affects neuronal migration patterns^72, 73^, and *PCLO* (ranked 4), which is important for synaptic vesicle trafficking and neurotransmitter release^74–76^. Although <u>predominant therapies for MDD target monoamine neurotransmitters, especially serotonin, the enrichment for GABAergic neurons is independent of serotonin pathways, suggesting that they might include new therapeutic targets for MDD.</u> These results were robustly detected in an independent brain scRNA-seq dataset, both in the gene programs (average correlation: 0.77 for the same cell type and -0.21 otherwise) and disease enrichments (0.77 correlation of the E-score over all cell type and trait pairs), including GABAergic neurons in MDD and BMI as well as glutamatergic neurons in insomnia and SCZ. Enrichment results for the remaining 49 diseases and traits in conjunction with brain cell type programs are reported in **Supplementary Fig. 9**.

Tissue specificity of both the cell type program and enhancer-gene strategy was important for successful linking, which we found by comparing the enrichment of all four possible combinations of immune or brain cell type programs with immune- or brain-specific enhancer-gene linking strategies, meta-analyzed across 11 immune-related diseases or 11 psychiatric/neurological diseases and traits (**Fig. 3d**). This highlights the importance of leveraging the tissue specificity of enhancer-gene strategies.

The 12 brain cellular process programs showed that the significant enrichment of brain-related diseases in neuronal cell types above is primarily driven by finer programs reflecting neuron subtypes (**Fig. 3f, Table 1, Supplemental Note**). For example, the enrichment of GABAergic neurons for BMI was driven by programs reflecting LAMP5^+^ and VIP^+^ subsets. Furthermore, the enrichment of GABAergic neurons for MDD reflects SST^+^ and PVALB^+^ subsets. We also observed enrichment in more specific cell subsets within glutamatergic neurons (e.g. IT neurons were enriched for neuroticism). Among inter cell type programs, electron transport cellular process programs (GABAergic and glutamatergic neurons) were enriched for several psychiatric/neurological traits, such as years of education, consistent with previous studies^77^.

### Linking cell types from diverse human tissues to disease

Analysis of kidney, liver, heart, skin and adipose cell types **(Supplementary Table 1)** and corresponding relevant traits (**Supplementary Table 2**) revealed the role of particular immune, stromal and epithelial cellular compartments across different diseases/traits. For example, kidney and liver cell type programs (**Supplementary Fig. 1**) highlighted relations with urine biomarker traits (**Fig. 4a, Supplementary Fig. 9 and 11a,b**), such as enrichment for creatinine level in kidney proximal and connecting tubule cell types, but not in liver cell types, as expected^78, 79^, or a significant enrichment for bilirubin level only in liver hepatocytes (driven by *ANGPTL3*; ranked 4)^80, 81^. In heart (**Fig. 4B, Supplementary Fig. 9 and 11c**, **Table 1**), atrial cardiomyocytes were enriched for atrial fibrillation, and pericyte and smooth muscle cells for blood pressure, consistent with their respective roles in determining heart rhythm through activity^82^ of ion channels (top genes included the ion channel genes *PKD2L2* (ranked 2), *CASQ2* (ranked 7) and *KCNN2* (ranked 18)) and blood pressure regulation through vascular tone^83^ (top genes driving included adrenergic pathway genes *PLCE1* (ranked 1), *CACNA1C* (ranked 21), and *PDE8A* (ranked 23)). In skin (**Fig. 4c, Supplementary Fig. 9**, **Table 1**), both BDNF signaling and Langerhans cells were enriched for eczema. Langerhans cells have been implicated in inflammatory skin processes related to eczema^84^ (top driving genes included IL-2 signaling pathway genes (*FCER1G* (ranked 3)*, NR4A2* (ranked 26), and *CD52* (ranked 43), which modulate eczema pathogenesis^85^). In adipose (**Fig. 4d, Supplementary Fig. 9 and 11e**), adipocytes were enriched for BMI, driven by adipogenesis pathway genes^86^ (*STAT5A* (ranked 15)*, EBF1* (ranked 29), *LIPE* (ranked 45) and triglyceride biosynthesis genes^86^ (*GPAM* (ranked 14), *LIPE* (ranked 45), both of which contribute to the increase in adipose tissue mass in obesity^87, 88^).

**Fig. 4.**
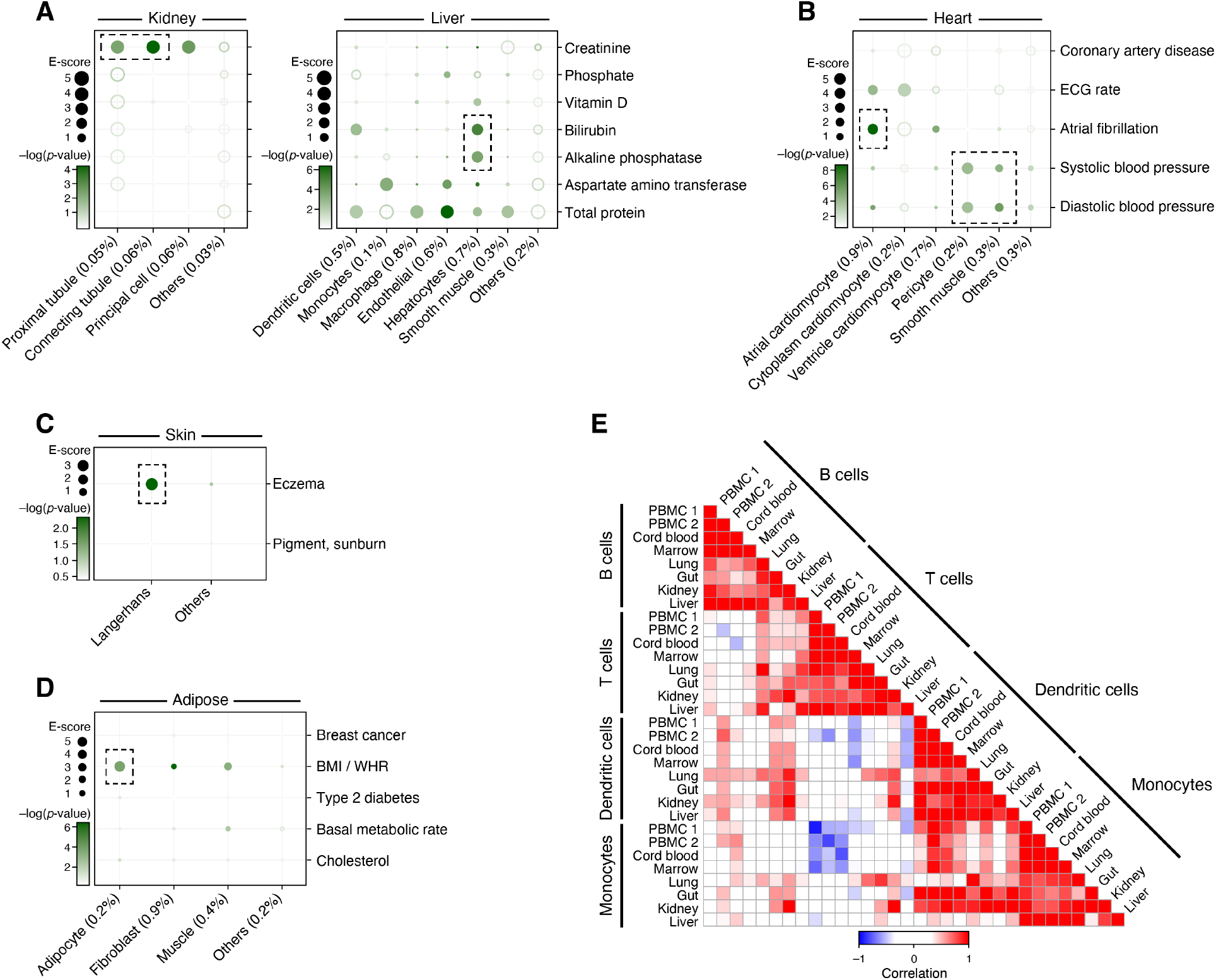
Linking cell types from diverse human tissues to disease. **a-d**. Enrichments of cell type programs for corresponding diseases and traits. Magnitude (E-score, dot size) and significance (-log_10_(P-value), dot color) of the heritability enrichment of cell type programs (columns) for diseases and traits relevant to the corresponding tissue (rows) for kidney and liver (a), heart (b), skin (c) and adipose (d). The size of each corresponding SNP annotation (% of SNPs) is reported in parentheses. Numerical results are reported in **data file S1**. Further details of all traits analyzed are provided in **Supplementary Table 2**. **e.** Correlation of immune cell type programs across tissues. Pearson correlation coefficients (color bar) of gene-level program memberships for immune cell type programs across different tissues (rows, columns), grouped by cell type (labels).

### Rare examples linking cell types from one tissue to disease manifestation in another tissue

We expanded our analysis to evaluate all cell type programs for all diseases/traits, irrespective of the tissue locus of disease aiming to identify cell type enrichments involving “mismatched” cell type -disease/trait pairs (**Supplementary Figure 5**). As expected, in most cases “mismatched” cell type programs and disease/trait pairs do not yield significant association. Notable exceptions included enrichments of skin Langerhans cells for Alzheimer’s disease (AD) (E-score: 15.2, p=10^-4^), M cells (in colon) for asthma (E-score: 2.2, p=10^-4^), and heart smooth muscle cells for lung capacity (E-score: 5.6, p=3*10^-4^).

In some cases, the association may indicate a direct relationship, whereas in other cases the associated cell type may only “tag” the causal cell type in the disease tissue, as cell type programs derived from cells of the same type across tissues were found to be highly correlated (**Fig. 4e**) with consistent enrichment in these correlated cell type programs (**Supplementary Fig. 5 and 9**). The enrichment of Langerhans cells for AD is plausible given that Langerhans cells respond differently to Aβ peptides, which has implications in AD immunotherapy^89^. On the other hand, the enrichment of colon M cells for asthma may suggest a role for lung-resident M cells, which have not been identified to date but are expected to be in the lung, as M cells stimulate IgA antibody production as an immune response^90^, while selective IgA immunodeficiency increases risk for asthma^91^. Similarly, the heart smooth muscle cell program may merely mirror that of airway smooth muscle cells, whose function is a pivotal determinant of lung capacity^92^.

### Linking neurons, microglia, and complement and apelin signaling pathways to MS and AD progression

We next turned to cases where both healthy and disease tissue have been profiled, allowing us to identify heritability in programs associated with disease-specific biology. Such understanding is especially important for identifying therapeutic targets associated with disease progression rather than disease onset mechanisms.

We first examined disease progression programs in multiple sclerosis (MS) and Alzheimer’s disease (AD) , where aberrant interactions between neurons and immune cells are thought to play an important role. We analyzed MS and AD GWAS data (**Supplementary Table 2**) along with cell type, disease progression, and cellular process programs from scRNA-seq of healthy and MS^49^ or AD^30^ brain (**Fig. 5a,e, Supplementary Table 1**). We considered brain enhancer-gene links (since MS and AD are neurological diseases), immune enhancer-gene links (since MS and AD are immune-related diseases) and non-tissue-specific enhancer-gene links (**Supplementary Fig. 12**) and detected strongest enrichment results for the immune enhancer-gene links. In both MS and AD, disease progression programs in each cell type differed substantially from cell type programs constructed from cells from healthy (r=0.16) or disease (r=0.29) samples alone (**Supplementary Fig. 13**). Furthermore, we confirmed that disease GWAS matched to the corresponding disease progression programs produced the strongest enrichments, although there was substantial cross-disease enrichment (**Supplementary Fig. 14**).

**Fig. 5.**
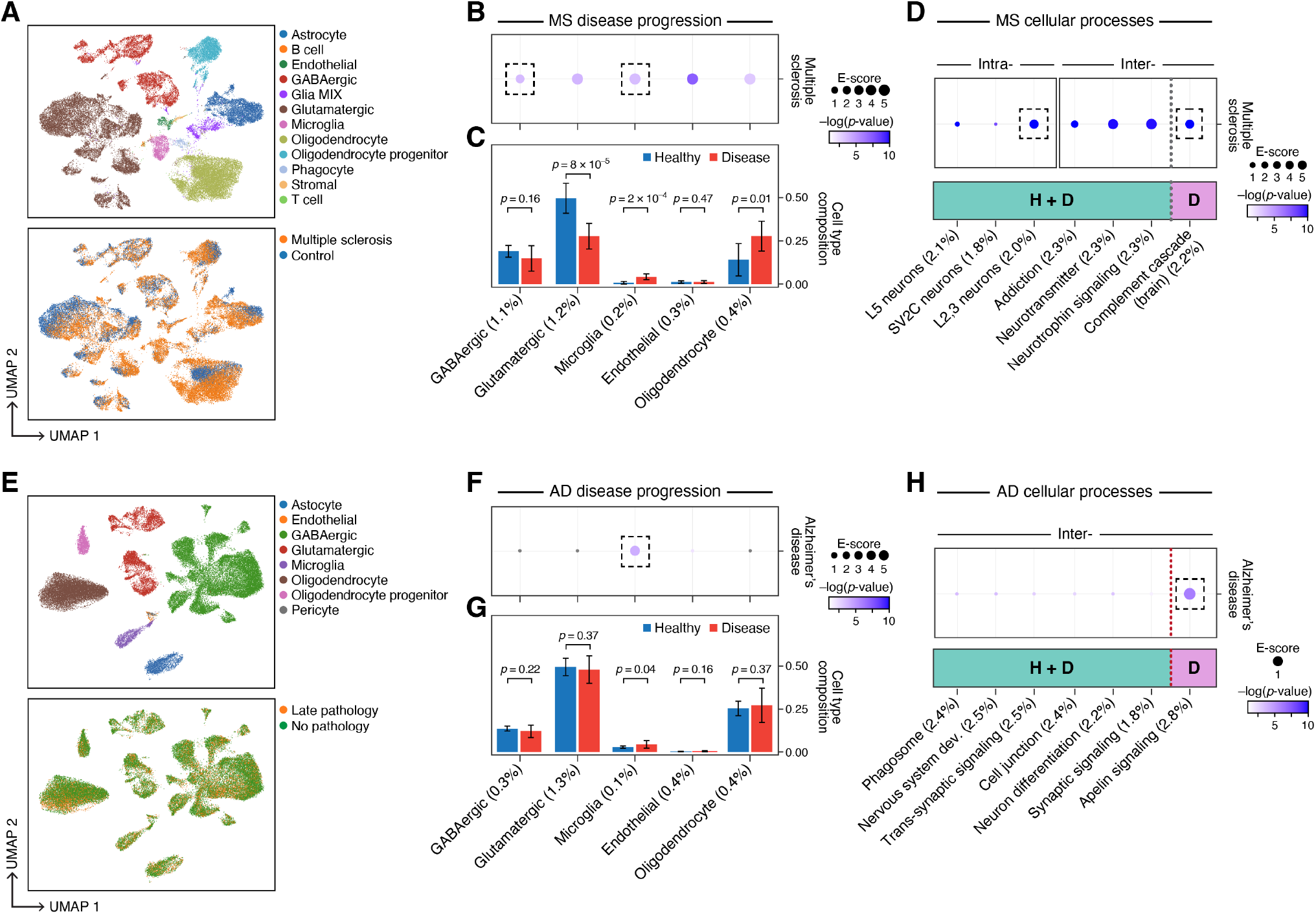
Linking MS and AD disease progression and cellular process programs to MS and AD. **a.** UMAP embedding of scRNA-seq profiles (dots) from MS and healthy brain tissue, colored by cell type annotations (top) or disease status (bottom). **b.** Enrichments of MS disease progression programs for MS. Magnitude (E-score, dot size) and significance (-log_10_(P-value), dot color) of the heritability enrichment of MS disease progression programs (columns), based on the Roadmap∪ABC-immune enhancer-gene linking strategy. **c.** Proportion (mean and SE) of the corresponding cell types (columns) in healthy (blue) and MS (red) brain samples. P-value: Fisher’s exact test. **d.** Enrichments of MS cellular process programs for MS. Magnitude (E-score, dot size) and significance (-log_10_(P-value), dot color) of the heritability enrichment of intra-cell type (left) or inter-cell type (right) cellular processes (healthy-specific (H), MS-specific (D) or shared (H+D)) (columns), based on the Roadmap∪ABC-immune enhancer-gene linking strategy. **e.** UMAP embedding of scRNA-seq profiles (dots) from AD and healthy brain tissue, colored by cell type annotations (top) or disease status (bottom). **f.** Enrichments of AD disease progression programs for AD. Magnitude (E-score, dot size) and significance (-log_10_(P-value), dot color) of the heritability enrichment of AD disease progression programs (columns), based on the Roadmap∪ABC-immune enhancer-gene linking strategy. **g.** Proportion (mean and SE) of the corresponding cell types (columns) in healthy (blue) and AD (red) brain samples. P-value: Fisher’s exact test. **h.** Enrichments of AD cellular process programs for AD. Magnitude (E-score, dot size) and significance (-log_10_(P-value), dot color) of the heritability enrichment of inter-cell type cellular processes (AD-specific (D) or shared (H+D)) (columns), based on the Roadmap∪ABC-immune enhancer-gene linking strategy. In panels b,c,d,f,g,h, the size of each corresponding SNP annotation (% of SNPs) is reported in parentheses. Numerical results are reported in **data file S2,3**. Further details of all traits analyzed are provided in **Supplementary Table 2**.

In MS, there was enrichment in disease progression programs in GABAergic neurons and microglia (**Fig. 5b**, **Supplementary Fig. 15**), as well as in Layer 2,3 glutamatergic neurons and the complement cascade (in multiple cell types) (**Fig. 5d**). The specific enrichment of the GABAergic neuron disease progression program (but not the healthy cell type program) for MS is consistent with the observation that inflammation inhibits GABA transmission in MS^94^. The GABAergic disease progression program was enriched with hydrogen ion transmembrane transporter activity genes, while the GABAergic cell type program was enriched in genes with general neuronal functions (**data file S9**). The enrichment of the microglia disease progression for MS is consistent with the role of microglia in inflammation and demyelination in MS lesions^95, 96^ and highlights a contribution of microglia in both disease onset and response. The top driving genes for the microglia disease progression enrichment included *MERTK* (ranked 2) and *TREM2* (ranked 4), both having roles in myelin destruction in MS patients^97, 98^. Supporting this finding, there is a significant increase in the number of microglia (p-value: 2x10^-4^, Fisher’s exact test) and a significant decrease in number of glutamatergic neurons (p-value: 8x10^-5^) in MS lesions (**Fig. 5c, data file S10**). In addition, there was enrichment for the complement cascade disease-specific cellular process program (in B cells and microglia; the top driving genes included FC-complement genes *CD37*, *FCRL2* and *FCRL1* (ranked 1, 10, 14) consistent with studies showing that Complement activity is a marker for MS progression^99–101^.

In AD, all associations highlighted the central role of microglia, suggesting that different processes may be at play at microglia or microglia subsets in healthy brain and after disease initiation: only the microglia disease progression program was enriched out of 8 disease progression programs tested (**Fig. 5e,f, Supplementary Fig. 16**), along with the healthy microglia program, and the apelin signaling pathway disease-specific cellular process program (inter cell type; GABAergic neurons and microglia). The microglia program enrichments are consistent with the contribution of microglia-mediated inflammation to AD progression^102, 103^. The top genes driving enrichment specifically in the disease progression program (but not the healthy cell type program) included *PICALM1, APOC1*, *APOE* and *TREM2* (ranked 1, 2, 3 and 8). *APOE* regulates microglial responses to Alzheimer’s related pathologies^104–106^, *APOC1* is a an *APOE*-dependent suppressor of glial activation^107^, and *TREM2* modulates microglial morphology and neuroinflammation in Alzheimer’s disease pathogenesis models^108, 109^. Supporting this finding, there is a significant increase in the number of microglia in AD brain (**Fig. 5g, data file S10**). The apelin signaling pathway disease-specific cellular process program is consistent with recent studies implicating this pathway in reducing neuroinflammation in animal models of Alzheimer’s disease^110, 111^. The top genes driving the enrichment included *SORL1* and *SYK* (ranked 2 and 3). *SORL1* expression levels are significantly reduced in Alzheimer’s disease patients, and has also been implicated by rare variant analyses^112–114^.

**Fig. 6.**
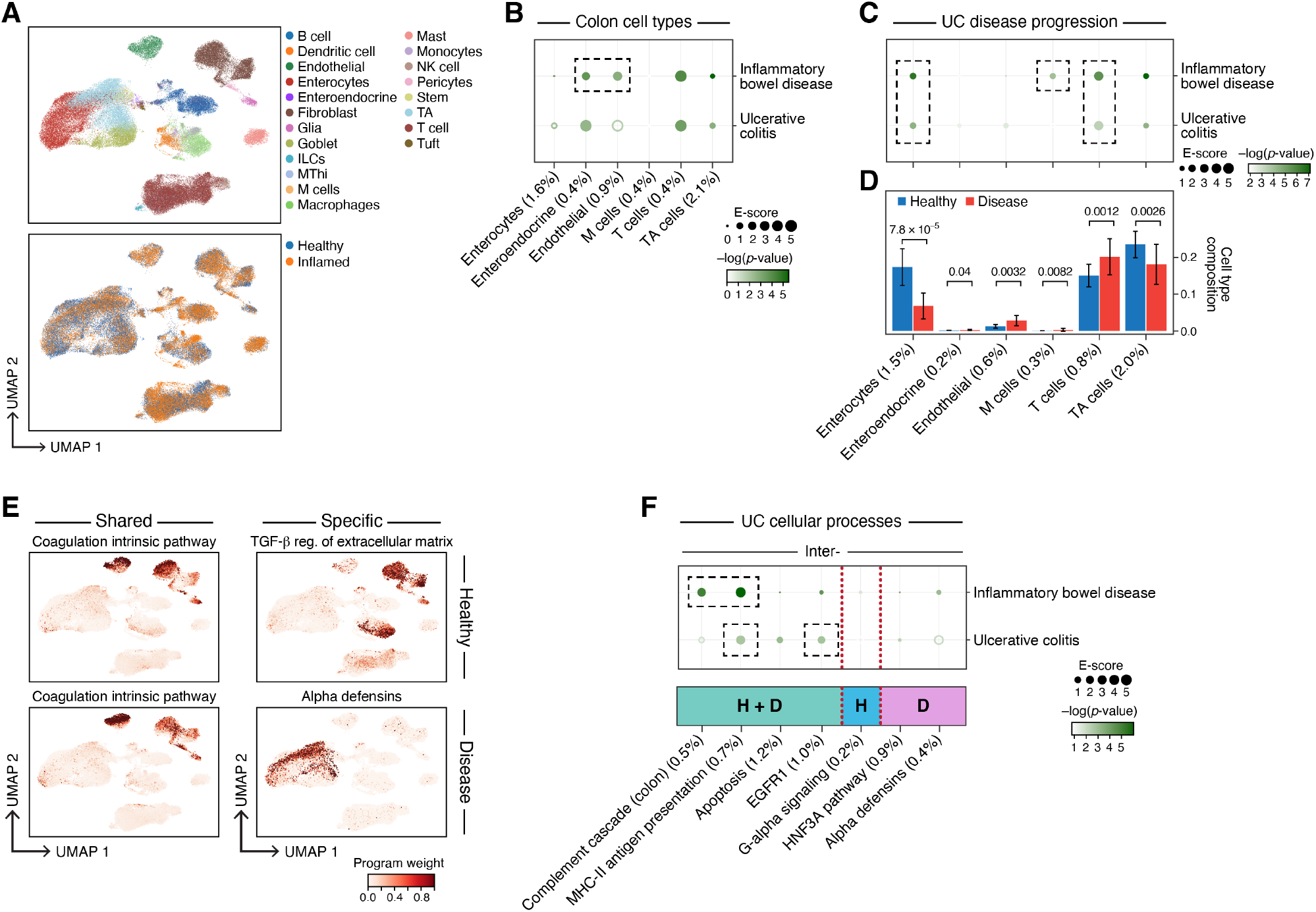
Linking UC disease progression and cellular process programs to UC and IBD. **a**. UMAP embedding of scRNA-seq profiles (dots) from UC and healthy colon tissue, colored by cell type annotations (top) or disease status (bottom). **b.** Enrichments of healthy colon cell types for disease. Magnitude (E-score, dot size) and significance (-log_10_(P-value), dot color) of the heritability enrichment of colon cell type programs (columns) for IBD or UC (rows). Results for additional cell types, including immune cell types in colon, are reported in **Supplementary Fig. 9 and data file S1**. **c.** Enrichments of UC disease progression programs for disease. Magnitude (E-score, dot size) and significance (-log_10_(P-value), dot color) of the heritability enrichment of UC disease progression programs (columns) for IBD or UC (rows). **d.** Proportion (mean and SE) of the corresponding cell types (columns) in healthy (blue) and UC (red) colon samples. P-value: Fisher’s exact test. **e.** Examples of shared (healthy and disease), healthy-specific, and disease-specific cellular process programs. UMAP (as in a), colored by each program weight (color bar) from NMF. **f.** Enrichments of UC cellular process programs for disease. Magnitude (E-score, dot size) and significance (-log_10_(P-value), dot color) of the heritability enrichment of inter-cell type cellular processes (shared (H+D), healthy-specific (H), or disease-specific (D)) (columns) for IBD or UC (rows). In panels b,c,d,f, the size of each corresponding SNP annotation (% of SNPs) is reported in parentheses. Numerical results are reported in **data file S1,2,3**. Further details of all traits analyzed are provided in **Supplementary Table 2**.

Thus, in both MS and AD, heritability was enriched in distinct ways in microglia cell type, disease progression and cellular process programs, suggesting new therapeutic opportunities to combat the role of microglia in varying contexts for disease risk and highlighting the importance of a multi-faceted analysis.

### Linking enterocytes and M cells to ulcerative colitis disease progression

We next examined the role of cell type, disease progression and cellular process programs in ulcerative colitis (UC), where failure to maintain the colon’s epithelial barrier results in chronic inflammation. We analyzed UC and IBD GWAS data (**Supplementary Table 2**) with healthy cell type, UC disease progression and UC cellular process programs constructed from scRNA-seq from healthy colon, and from matched uninflamed and inflamed colon of UC patients (**Fig. 6a, Supplementary Table 1**). We compared colon enhancer-gene links (**Fig. 6**) and non-tissue-specific enhancer-gene links (**Supplementary Fig. 12**) and detected strongest enrichment results for the colon enhancer-gene links. As in MS and AD, UC disease progression programs in each cell type differed substantially from corresponding healthy or disease colon cell type programs (average Pearson *r*=0.24; **Supplementary Fig. 13, data file S11**).

In addition to previously observed enrichments in healthy immune cell type programs, our analysis highlighted healthy cell type programs of enteroendocrine and endothelial cells, disease progression programs of enterocytes and M cells, as well as the complement cascade (in plasma, B cells, enterocytes and fibroblasts), MHC-II antigen presentation (macrophages, monocytes and dendritic cells), and EGFR1 signaling (macrophages and enterocytes) in both healthy and disease cells (**Fig. 6, Supplementary Fig. 9**, **data file S1**). The strong enrichment in endothelial cells, which comprise the gut vascular barrier, is consistent with their rapid changes in UC^115^; the top driving genes included members of the TNF-" signaling pathway (*EFNA1*, *NFKBIA*, *CD40*, ranked 18, 26, 29), a key pathway in UC^116^.

The disease progression programs (**Fig. 6c, Table 1, Supplementary Fig. 15 and 11**) highlighted M cells, a rare cell type in healthy colon that increases in UC^34^ (**Fig. 6d**, **data file S10**). M cells surveil the lumen for pathogens and play a key role in immune–microbiome homeostasis^117^. Supporting this finding, mutations in *FERMT1*, a top driving gene in the M cell disease progression program (ranked 3), cause Kindler syndrome, a monogenic form of IBD with UC-like symptoms^118–120^. Notably, there was no enrichment in M cell healthy cell type programs (**Fig. 6b**), emphasizing that M cells are activated specifically in UC disease, as their proportions increase (p=0.008) **(Fig. 6d**).

### Immune and connective tissue cell types linked to asthma disease progression

We analyzed GWAS data for asthma, IPF, COVID-19 (both general COVID-19 and severe COVID-19), and lung capacity (**Supplementary Table 2**) with healthy cell type, disease progression and cellular process programs from asthma, IPF, COVID-19 and healthy^29^ (lower lung lobes) tissue scRNA-seq (**Fig. 7a,c,f**, **Supplementary Fig. 13d-f and 15, data file S11**), using either lung enhancer or immune enhancer gene links. For asthma, there was significant enrichment for healthy cell type and disease progression programs in T cells (see **Supplemental Note**), and for lung capacity (height-adjusted FEV1adjRVC), there was significant enrichment for healthy cell type and disease progression programs in fibroblasts (**Fig. 7b**, **data file S1**) and the MAPK cellular process program (in basal, club, fibroblast and endothelial cells) (**Fig. 7f, g**, **Table 1**). For IPF and COVID-19, the enrichment results are detailed in the **Supplemental Note**.

**Fig. 7.**
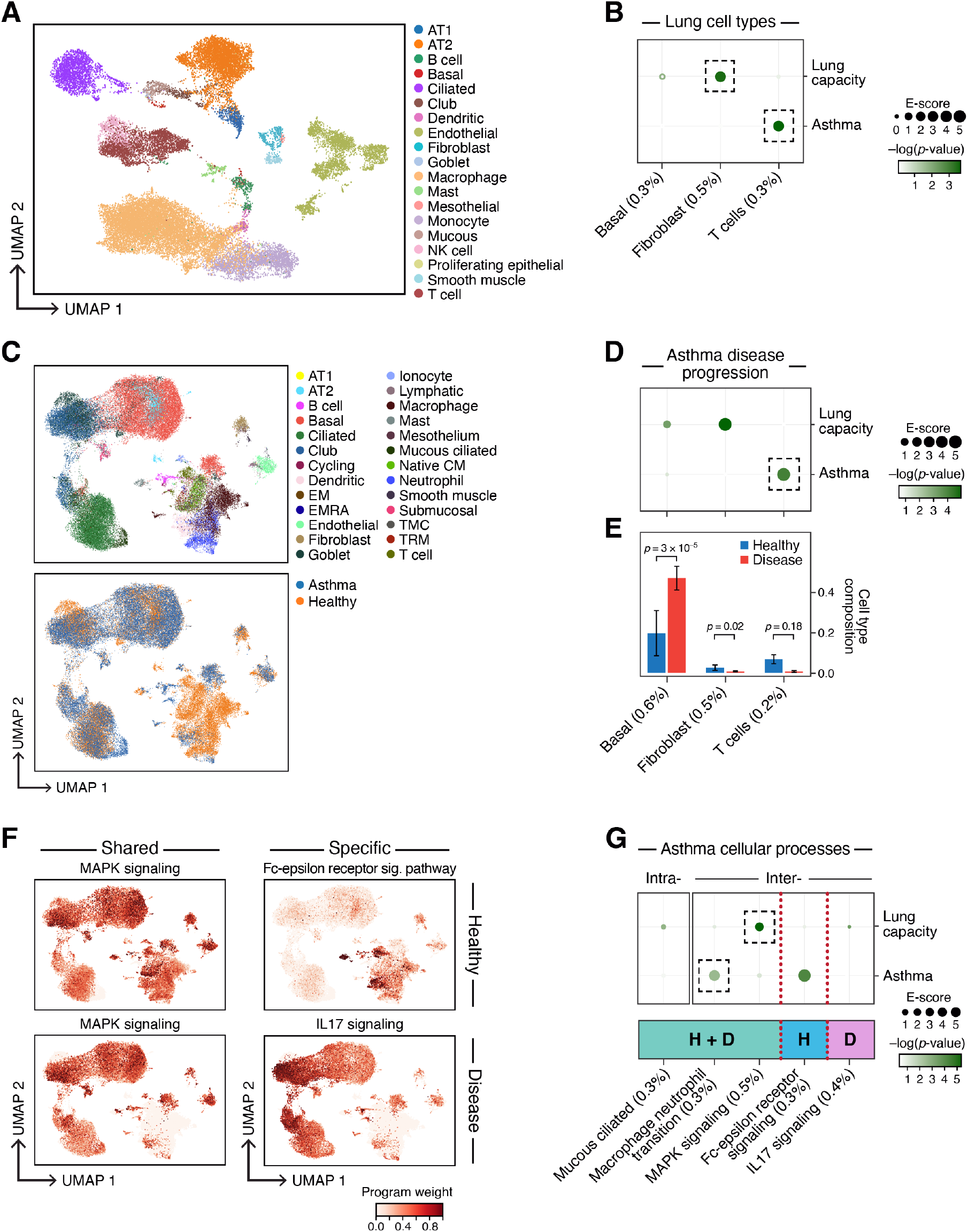
Linking asthma disease progression and cellular process programs to asthma and lung capacity. **a**. UMAP embedding of healthy lung scRNA-seq profiles (dots) colored by cell type annotations. **b.** Enrichments of healthy lung cell types for disease. Magnitude (E-score, dot size) and significance (-log_10_(P-value), dot color) of the heritability enrichment of healthy lung cell type programs (columns) for lung capacity or asthma (rows). **c.** UMAP embedding of scRNA-seq profiles (dots) from asthma and healthy lung tissue, colored by cell type annotations (top) or disease status (bottom). **d.** Enrichments of asthma disease progression programs for disease. Magnitude (E-score, dot size) and significance (-log_10_(P-value), dot color) of the heritability enrichment of asthma disease progression programs (columns) for lung capacity or asthma (rows). **e.** Proportion (mean and SE) of the corresponding cell types (columns), in healthy (blue) and asthma (red) lung samples. P-value: Fisher’s exact test. **f.** Examples of shared (healthy and disease), healthy-specific, and disease-specific cellular process programs. UMAP (as in c), colored by each program weight (color bar) from NMF. **g.** Enrichments of asthma cellular process programs for disease. Magnitude (E-score, dot size) and significance (-log_10_(P-value), dot color) of the heritability enrichment of intra-cell type (left) and inter-cell type (right) cellular processes (shared (H+D), healthy-specific (H), or disease-specific (D)) (columns) for lung capacity and asthma GWAS summary statistics (rows). In panels b,d,e,g, the size of each corresponding SNP annotation (% of SNPs) is reported in parentheses. Numerical results are reported in **data file S1,2,3**. Further details of all traits analyzed are provided in **Supplementary Table 2.**

For example, both healthy and disease progression fibroblast/stromal programs were enriched for lung capacity (but not asthma), consistent with the adverse impact of overproduction of extracellular matrix (ECM) on the reduced lung capacity and elasticity characteristic of fibrosis^121^. In the cell type program, top driving genes included *LOX* (ranked 1), which alters ECM mechanical properties via collagen cross-linking^122^, and *TGFBR3* (ranked 37) which regulates the pool of available TGFβ, a master regulator of lung fibrosis. Notably, the enrichment of basal cell disease progression programs, but not healthy cell type programs, in lung capacity are supported by the significant increase (p-value: 3x10^-5^) in basal cells in asthma *vs*. healthy lungs (**Fig. 7e**). Expanding the analysis to cellular process programs, the top driving genes of the enrichment of a MAPK signaling pathway program for lung capacity (in basal, club, fibroblast and endothelial), include *FOXA3* (ranked 1), which plays a key role in allergic airway inflammation^123^, and *PDE2A* (ranked 2), which has been associated with alveolar inflammation^124^.

## DISCUSSION

Prior work on identifying disease-critical tissues and cell types by combining expression profiles and human genetics signals has largely focused on the direct mapping of the expression of individual genes^34^ and genome-wide polygenic signals^18, 36^ to discrete cell categories. Our study demonstrates that there is much to be gained by linking inferred representations of the underlying biological processes beyond cell types in different cell and tissue contexts with genome-wide polygenic disease signals, by integrating scRNA-seq, epigenomic and GWAS data sets.

Our work introduces three main conceptual advances. First, by integrating scRNA-seq data and GWAS summary statistics using tissue-specific enhancer-gene linking strategies, we detect subtle differences in SNP to gene mapping between tissues which upon aggregation over the full GWAS signal produce strong differences in disease heritability across cell types. Second, by constructing disease progression programs comparing cells of the same type in disease *vs*. healthy tissue, we project GWAS signals across disease-specific cell states. Third, by using NMF to construct cellular process programs that do not rely on known cell type categories, we identify cellular mechanisms that vary across a continuum of cells of one type or are shared between cells of different types such as the MAPK signaling pathway identified in the lung.

Leveraging these advances, we identified notable enrichments (**Table 1**) that have not previously been identified using GWAS data and are biologically plausible but not clearly expected, thus providing important new knowledge. We also observed patterns across datasets that offer new insights. For example, we observed that disease progression programs, but not healthy cell type programs, of epithelial cells (M cells and basal cells) tend to be enriched in autoimmune diseases (UC and asthma). In contrast, for immune cells healthy and disease progression programs tended to be similarly enriched. We posit that this suggests a role for epithelial cells in disease progression over initiation. Future studies are required to experimentally validate these new hypotheses.

Our work has several limitations that highlight directions for future research. First, the enhancer-gene linking strategies from Roadmap and Activity-By-Contact (ABC) models are limited in the tissues and cell states represented. More fine-grained enhancer-gene linking strategies will likely prove beneficial, but the strategies that we used here provide a clear improvement over a standard gene window-based approach. Second, we focus on genome-wide disease heritability (rather than a particular locus); however, our approach can be used to implicate specific genes and gene programs. Third, sc-linker does not distinguish whether two cell types (or more generally, gene programs) implicated in disease exhibit conditionally independent signals. Assessing this via a conditional S-LDSC analysis of the corresponding SNP annotations is likely to be underpowered, as the gene programs (and SNP annotations) may be highly correlated. A more powerful approach may be to define cell type programs based on specific expression relative to a narrower set of cells. Fourth, although all studies considered in this work profiled large numbers of cells (up to 300,000 in some tissues), some rare cell types and processes may not yet be adequately sampled due to the number of cells or their tissue distribution^125^, or may only be apparent in a disease context, as we observe for rare M cells in UC. Fifth, we have focused on human scRNA-seq data^33^; however, incorporating data from animal models, as discussed in prior work^36^, would allow experimental validation of disease mechanisms in model organisms. Sixth, the disease progression programs that we link to disease may not be causal for disease, but rather reflect disease-induced changes or genetic susceptibility to disease^126, 127^. However, our findings clearly validate the relevance of these gene programs to disease as observed in M cells and UC^34^. Seventh, the LD score regression framework^11^ is primarily applicable to common and low-frequency variants, and less applicable to rare variant enrichments. Eighth, we capture programs by cell category or gene co-variation, whereas future work could extend beyond these to capture dynamic cellular transitions^128^.

Looking forward, the gene program-disease links identified by our analyses can be used to guide downstream studies, including designing systematic perturbation experiments^129, 130^ in cell and animal models^131^ for functional follow up. We anticipate that gene programs will continue to grow and refine due to the continued growth of different types of profiling data – including from single-cell atlases across many tissues and diseases, Perturb-seq^129^ experiments under genetic or chemical perturbation, spatial transcriptomics, and other modalities^132, 133^. Such analyses can then expand to genetic interactions within and between cells. In the long term, with the increasing success of PheWAS and the integration of multi modal single cell resolution epigenomics, this framework will continue to be useful in identifying biological mechanisms driving a broad range of diseases.

## METHODS

### scRNA-seq data pre-processing

All scRNA-seq datasets in this study^25–30, 34, 45–51^ are publicly available cell by gene expression matrices that are aligned to the hg38 human transcriptome (**Supplementary Table 1**). Each dataset included metadata information for each cell describing the total number of reads in the cell and which sample the cell corresponds to and, if applicable, its disease status. We transformed each expression matrix to a count matrix by reversing any log normalization processing (because each downloaded dataset contained either (i) raw counts, (ii) normalized log_2_ TP10K, or (iii) normalized log_10_ TP10K), and standardized the normalization approach across all datasets to account for differences in sequencing depth across cells by normalizing by the total number of UMIs per cell, converting to transcripts-per-10,000 (TP10K) and taking the log of the result to obtain log(10,000*UMIs/total UMIs + 1) “log_2_(TP10K+1)” as the final expression unit.

### Dimensionality reduction, batch correction, clustering and annotation of scRNA-seq

The log_2_(TP10K+1) expression matrix for each dataset was used for the following downstream analyses. For each dataset, we identified the top 2,000 highly variable genes across the entire dataset using Scanpy’s^42^ *highly_variable_genes* function with the sample ID as input for the batch. We then performed a Principal Component Analysis (PCA) with the top 2,000 highly variable genes and identified the top 40 principle components (PCs), beyond which negligible additional variance was explained in the data (the analysis was performed with 30, 40, and 50 PCs and was robust to this choice). We used Harmony^134^ for batch correction, where each sample was considered its own batch. Subsequently, we built a *k*-nearest neighbors graph of cell profiles (*k* = 10) based on the top 40 batch corrected components computed by Harmony and performed community detection on this neighborhood graph using the Leiden graph clustering method^135^ with resolution 1. For each dataset, individual single-cell profiles were visualized using the Uniform Manifold Approximation and Projection (UMAP)^136^. If prior annotations were available they are used as a reference to annotate each cell in each dataset. If prior annotations were not available, we used established cell type-specific expression signatures and gene markers described in the data source to annotate cells at the resolution of Leiden clusters.

### Cell type gene programs

We constructed cell type programs for every cell type in a given tissue by applying a non-parametric Wilcoxon rank sum test for differential expression (DE) between each cell type *vs*. other cell types and computed a p value for each gene. Using a previously published strategy^15^, we transform these p-values to X = -2 log (p), which follow a 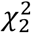 distribution, and these transformed values to a grade between 0 and 1 using the min max normalization g = ( X – min (X) )/( max(X) – min(X) ) resulting in a relative weighting of genes in each program. In brief, cell type programs constructed from healthy cells were termed as healthy cell type programs and similarly cell type programs constructed from disease cells were termed as disease cell type programs.

### Disease progression gene programs

We constructed disease progression programs for each cell type observed in both healthy and matching disease tissue. For each cell type, we computed a gene-level non-parametric Wilcoxon rank sum DE test between cells from healthy and disease tissues of the same cell type. The p-values for each gene were transformed to a grade between 0 and 1 using the same strategy as in the cell type program to form a relative weighting of genes in each program. In the COVID-19 BAL scRNA-seq, we also constructed viral progression programs based on differential expression between viral infected and uninfected cells of the same cell type in COVID-19 disease individuals.

We observed low correlation between healthy cell type gene programs and disease progression gene programs (see **Supplementary Fig. 13 and data file S11**).

We observed low correlation between healthy cell type gene programs and disease progression gene programs (see **Supplementary Fig. 13 and data file S11**).

### Cellular process gene programs

Using latent factors derived from non-negative matrix factorization (NMF)^43^ (see below), we define a cellular process program based on genes with high correlation (across cells) between their expression in each cell and the contribution of the factor to each cell (collapsing latent factors with high correlation). The correlations were transformed to a probabilistic scale (between 0 and 1) by scaling their values (negative correlations are assigned to 0). We then annotated each factor (program) by the pathway most enriched in the top driving genes for the factor and labeled each as an ‘intra-cell type’ or ‘inter-cell type’ latent factor if the pathway was highly correlated with only one or multiple cell type programs, respectively.

We constructed cellular process programs using an unsupervised approach, by applying non-negative matrix factorization (NMF)^43^ to the scRNA-seq cells-by-genes matrix. The solution to this formulation can be identified by solving the following minimization problem:

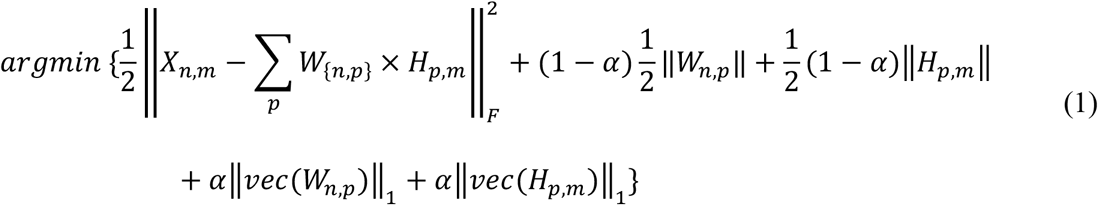

where *X_n,m_* represents the log-normalized expression of gene *m* in sample *n*, *W_n,p_* denotes the grade of membership of latent factor p in sample n, and *H_p,m_* represents the factor weight of factor *p* in gene *m.* NMF identifies cellular processes as latent factors with a grade of contribution to each cell. For each dataset, we specified the number of latent factors *p* to be the number of annotated cell types in the dataset plus 10. For each latent factor, we define a cellular process gene program by identifying genes with high correlation (across cells) between expression in a cell and the contribution of each factor to each cell. Latent factors with correlation above 0.8 are collapsed to only consider a single latent factor. We annotated each cellular process program by the pathway most enriched in the genes with highest correlation (across cells) between expression levels and factor weights (*H*) underlying the cellular process program (not necessarily the most highly expressed genes, **Supplementary Fig. 17**) and labeled it as an ‘intra-cell type’ or ‘inter-cell type’ cellular process program if highly correlated with only one or multiple cell type programs, respectively.

### Cellular process gene programs constructed from healthy and disease tissues

For scRNA-seq from healthy and disease tissue contexts, we propose a modified NMF approach to construct gene programs that are either shared across both tissues, specific to healthy tissue or specific to disease tissue. Let *H_p*N1_* be the observed gene expression data for a tissue *T* from a healthy individual and *D_p*N2_* be the observed gene expression data for the corresponding tissue from a disease individual. *P* is the number of features (genes) and *N*_1_ and *N*_1_ denote the number of samples from the healthy and disease tissues, respectively.

We assume a non-negative matrix factorization for 4 andas follows

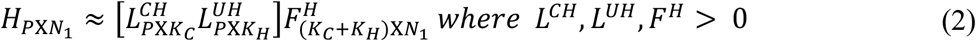

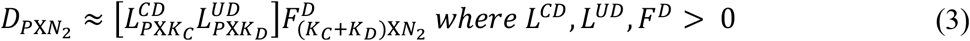

where *K_c_* is the number of shared programs between the healthy and the disease samples, *K_H_* is the number of healthy specific programs and *K_D_* is the number of disease-specific programs. *L^CH^* and *L^CD^* are used to denote the shared programs between healthy and disease states. Therefore, we assume that *L^CH^* is very close to *L^CD^* but not exact to account for other factors like experimental conditions perturbing the estimates slightly. On the other hand, *L^UH^* and *L^UD^* are used to denote the healthy-specific and disease-specific programs respectively. *F^H^* and *F^D^* denote the program weights in the healthy and disease samples respectively. frame this in the form of the following optimization problem

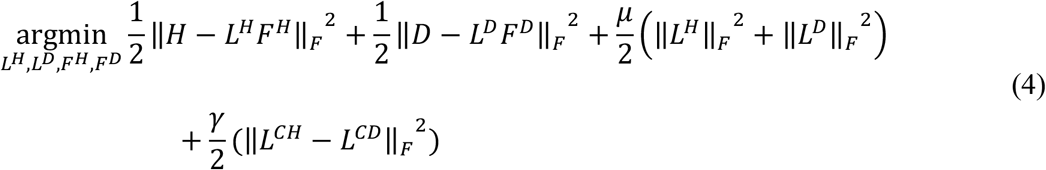

Where 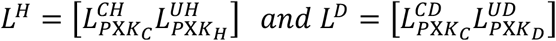 and γ is a tuning parameter that controls how close *L^CH^* is to *L^CD^*. *μ* represents a tuning parameter that controls for the size of the loadings and the factors.

To determine the multiplicative updates of the NMF optimization problem in Equation 4 we compute the derivatives of the optimization criterion with respect to each parameter of interest.

We call the optimization criterion as Q:

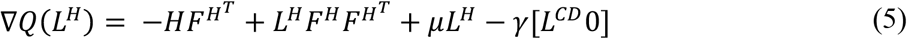

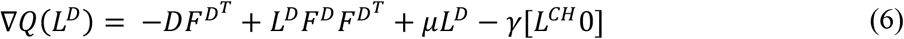

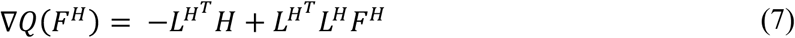

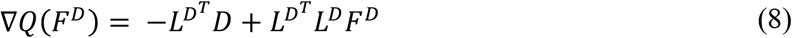

Following the multiplicative update rules of NMF as per Lee and Seung (NIPS 2001), we get the following iterative updates and assume convergence has been achieved after 100 iterations or when the reconstruction error is below a user-specified error threshold (here the threshold is taken to be 1e-04).

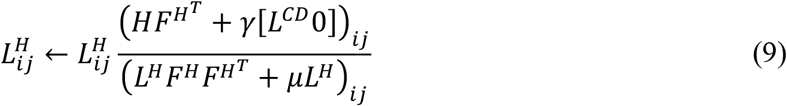

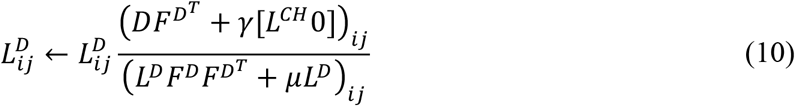

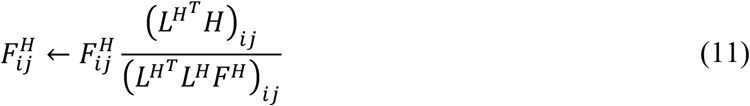

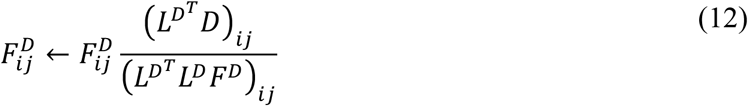

### Enhancer-gene linking strategies

We define an enhancer-gene linking strategy as an assignment of 0, 1 or more genes to each SNP with a minor allele count >5 in the 1000 Genomes Project European reference panel^137^. Here, we primarily considered an enhancer-gene linking strategy defined by the union of the Roadmap^21, 138^ and Activity-By-Contact (ABC)^22, 23^ strategies. Roadmap and ABC enhancer gene links are publicly available for a broad set of tissues and have been shown to outperform other enhancer-gene linking strategies in previous work^139^. We consider tissue-specific Roadmap and ABC enhancer-gene linking strategies for gene programs corresponding to any of the biosamples (cell types or tissues) associated with the relevant tissue. Based on analysis in immune cell types, 87% of genes expressed in the scRNA-seq were observed to have enhancer-gene links. We also consider non-tissue specific Roadmap and ABC strategies (**Supplementary Fig. 12**). Besides this enhancer-gene linking strategy, we also considered a standard 100kb window-based strategy^13, 18^.

### Genomic annotations and the baseline-LD models

We define an annotation as an assignment of a numeric value to each SNP in a predefined reference panel (*e.g.*, 1000 Genomes Project^137^; see Data Availability). Binary annotations can have value 0 or 1 only; continuous-valued annotations can have any real value; our focus is on continuous-valued annotations with values between 0 and 1. Annotations that correspond to known or predicted functions are referred to as functional annotations. The baseline-LD model^40, 41^ (v.2.1) contains 86 functional annotations (see Data Availability), including binary coding, conserved, and regulatory annotations (*e.g.*, promoter, enhancer, histone marks, TFBS) and continuous-valued linkage disequilibrium (LD)-related annotations.

### Stratified LD score regression

Stratified LD score regression (S-LDSC) assesses the contribution of a genomic annotation to disease and complex trait heritability^11^. S-LDSC assumes that the per-SNP heritability or variance of effect size (of standardized genotype on trait) of each SNP is equal to the linear contribution of each annotation.

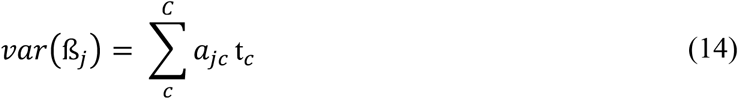

where *a_jc_* is the value of annotation *c* at SNP *j*, with the annotation either continuous or binary (0/1), and *t_c_* is the contribution of annotation *c* to per SNP heritability conditional on the other annotations. S-LDSC estimates t_c_ for each annotation using the following equation:

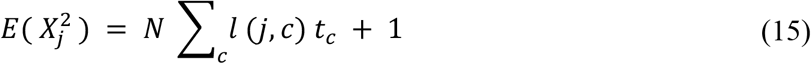

where 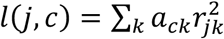 is the stratified LD score of SNP *j* with respect to annotation *c*, *r_jk_* is the genotypic correlation between SNPs *j* and *k* computed using 1000 Genomes Project, and *N* is the GWAS sample size.

We assess the informativeness of an annotation *c* using two metrics. The first metric is Enrichment score (E-score), which relies on the enrichment of annotation *c* (*E_c_* ), defined for binary annotations as follows (for binary and probabilistic annotations only):

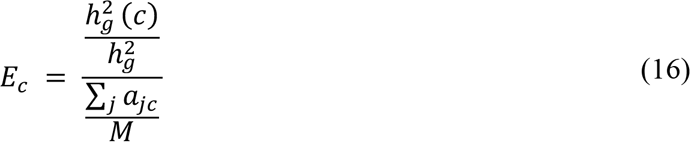

where 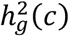 is the heritability explained by the SNPs in annotation *c,* weighted by the annotation values where *M* is the total number of SNPs on which this heritability is computed (5,961,159 in our analyses). The Enrichment score (E-score) is defined as the difference between the enrichment for annotation *c* corresponding to a particular program against a SNP annotation for all protein coding genes with a predicted enhancer-gene link in the relevant tissue. The E-score metric generalizes to probabilistic annotations with values between 0 and 1^44^. We primarily focus on the p-value for nonzero enrichment score (see below).

The second metric is standardized effect size (*τ*^∗^), the proportionate change in per-SNP heritability associated with a one standard deviation increase in the value of the annotation, conditional on other annotations included in the model^40^.

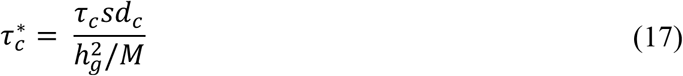

where *sd_c_* is the standard error of annotation *c*, 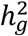 is the total SNP heritability and *M* is as defined previously. 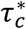 is the proportionate change in per-SNP heritability associated with an increase of one standard deviation in the value of a annotation.

We assessed the statistical significance of the enrichment score and *τ*^∗^ via block-jackknife, as in previous work^11^, with significance thresholds determined via False Discovery Rate (FDR) correction (q-value < 0.05)^140^. FDR was calculated over all relevant relatively independent traits for a tissue and all programs of a particular type (cell type programs, disease progression programs, cellular process programs) derived from that tissue. We used the p-value for nonzero enrichment score as our primary metric, because τ^∗^ is often non-significant for small cell-type-specific annotations when conditioned on the baseline-LD model^141^.

### GWAS summary statistics

We analyzed publicly available GWAS summary statistics for 60 unique diseases and traits with genetic correlation less than 0.9. Each trait passed the filter of being well powered enough for heritability studies (z score for observed heritability > 5). We used the summary statistics for SNPs with minor allele count >5 in a 1000 Genomes Project European reference panel^137^. The lung FEV1FVC trait was corrected for height data. For COVID-19, we analyzed two phenotypes – general COVID-19 (covid *vs*. population, liability scale heritability ℎ^2^ = 0.05, se. = 0.01), and severe COVID-19 (hospitalized covid vs population, liability scale heritability ℎ^2^ = 0.03, se. = 0.01)^142^ (meta-analysis round 4, October 20, 2020, https://www.covid19hg.org/).

### Computing a sensitivity/specificity index

We define a sensitivity/specificity index to benchmark (i) sc-linker vs. MAGMA gene-set enrichment analysis, and (ii) different versions of sc-linker corresponding to varying ways to define cell type programs and SNP-to-gene linking strategies

For the comparison of sc-linker with MAGMA, we define the sensitivity/specificity index as the difference of (i) the average of -log_10_(P-values) of enrichment score (association) using sc-linker (MAGMA) for “putatively positive control” (gene program, trait) combinations and (ii) the average of -log_10_(P-values) of gene-set level enrichment score (association) using sc-linker (MAGMA) for “putatively negative control” (gene program, trait) combinations. In **Fig. 4e**, the putatively positive control combinations include immune programs for blood cell traits and immune diseases, and brain programs for brain related traits; all other combinations are considered to be putatively negative controls. In **Supplementary Fig. 8**, the putatively positive control combinations include B and T cells for lymphocyte percentage, monocytes for monocyte percentage, megakaryocytes for platelet count, erythroid for RBC count and RBC distribution width; all other combinations of cell types and traits are considered as putatively negative controls.

For the comparison of the different versions of the sc-linker approach using either varying definitions of cell type programs (**Supplementary Fig. 6** and **7**) or different ways to link SNPs to genes beyond Roadmap∪ABC enhancer-gene linking strategy (**Fig. 3d,e** and **Supplementary Fig. 3**), we use a slightly different definition of sensitivity/specificity index. Instead of the -log P-value, we use the τ* metric from the S-LDSC method, which evaluates conditional information in the SNP annotation corresponding to a gene program, corrected for the annotation size. This metric is preferred when comparing across cell-type programs or enhancer-gene linking strategies that are widely different in their corresponding SNP annotation sizes, as is the case in these comparisons (we note that use of this metric is not possible in comparisons involving MAGMA, which does not estimate τ*).

### Identifying genes driving heritability enrichment

For each gene program, we first subset the full gene list to only consider genes with greater than 80% probability grade of membership in the gene program. Subsequently, we ranked all remaining genes using MAGMA (v 1.08) gene level significance score and considered the top 50 ranked genes for further downstream analysis, which is different from the top 200 genes used for a “baseline” method for scoring cell type enrichments for disease that we used as a benchmark for sc-linker.

### Identifying statistically significant differences in cell type proportions

To identify changes in cell type proportions between healthy and disease tissue, we used a multinomial regression test to jointly test changes across all cell types simultaneously. This helps account for all cell type changes simultaneously, as an increase in the number of cells of one cell types implies fewer cells of the other cell type will be captured. This regression model and the associated p-values were calculated using the multinom function in the nnet R package.

## DATA AVAILABILITY

All postprocessed scRNA-seq data (except for Alzheimer’s disease; see below), gene programs, enhancer-gene linking annotations, supplementary data files and high-resolution figures are publicly available online at https://data.broadinstitute.org/alkesgroup/LDSCORE/Jagadeesh_Dey_sclinker. The Alzheimer’s disease scRNA-seq data^30^ is available exclusively at https://www.radc.rush.edu/docs/omics.htm per its data usage terms. This work used summary statistics from the UK Biobank study (http://www.ukbiobank.ac.uk/). The summary statistics for UK Biobank used in this paper are available at https://data.broadinstitute.org/alkesgroup/UKBB/. The 1000 Genomes Project Phase 3 data are available at ftp://ftp.1000genomes.ebi.ac.uk/vol1/ftp/release/2013050. The baseline-LD annotations are available at https://data.broadinstitute.org/alkesgroup/LDSCORE/. We provide a web interface to visualize the enrichment results for different programs used in our analysis at: https://share.streamlit.io/karthikj89/scgenetics/www/scgwas.py.

## CODE AVAILABILITY

This work uses the S-LDSC software (https://github.com/bulik/ldsc) as well as MAGMA v1.08 for *post-hoc* analysis (https://ctg.cncr.nl/software/magma). Code for constructing cell type, disease progression and cellular process gene programs from scRNA-seq data and performing the healthy and disease shared NMF can be found at https://github.com/karthikj89/scgenetics. Code for processing gene programs and combining with enhancer-gene links can be found at https://github.com/kkdey/GSSG.

## ACKNOWLEDGMENTS

We thank Leslie Gaffney for assistance with preparing figures as well as Sijia Chen, Chris Smillie, Basak Eraslan, Alok Jaiswal, and the entire Price and Regev groups for helpful scientific discussions. **Funding:** This work was funded through (K.A.J) NIH F32 Fellowship, (A.L.P) NIH grants U01 HG009379, R01 MH101244, R37 MH107649, R01 MH115676 and R01 MH109978, and (A.R.) Klarman Cell Observatory, HHMI, the Manton Foundation and NIH grant 5U24AI118672.

## AUTHOR CONTRIBUTIONS

K.A.J., K.K.D, A.L.P and A.R designed the study. K.A.J., K.K.D. developed statistical methodologies and performed all computational analyses. A.L.P and A.R. provided expert guidance and feedback on analysis and results. D.T.M interpreted biological signals and guided K.A.J. and K.K.D. on highlighting biological insights. J.M.E. provided Activity-by-Contact mappings. S.G. provided guidance on enhancer-gene linking strategies. R.J.X. provided guidance on biological interpretations. K.A.J., K.K.D, A.L.P and A.R wrote the manuscript with detailed input from D.T.M. and feedback from all authors.

## EXTENDED DATA FILE LEGENDS

**Data File S1: Healthy cell type program heritability enrichment results.** Numerical values for E-score and significance are reported for all cell type programs and traits analyzed.

**Data File S2: Disease progression program heritability enrichment results.** Numerical values for E-score and significance are reported for all disease progression programs and traits analyzed.

**Data File S3: Cellular process program heritability enrichment results.** Numerical values for E-score and significance are reported for all healthy, disease, and shared cellular processes and traits analyzed.

**Data File S4: List of genes driving each enrichment.** Up to 50 genes with the strongest MAGMA gene score and membership in the gene program.

**Data File S5: Heritability enrichment results from eQTL, PCHi-C and other alternative enhancer-gene linking strategies.** Numerical values for E-score and significance are reported for all traits analyzed with alternative enhancer-gene linking strategies.

**Data File S6: Heritability enrichment results from alternative approaches for constructing cell type gene programs.** Numerical values for E-score and significance are reported for all traits analyzed with the alternative cell type programs.

**Data File S7: FUMA enrichments for blood cell traits and immune cell type programs.** Numerical values for beta, standard error and p-value for all cell types and traits analyzed.

**Data File S8: MAGMA gene set enrichment results for all cell type programs.** MAGMA scores across all traits analyzed.

**Data File S9: Pathway enrichment analysis for each disease progression program.** Gene overlap, p-value and gene list for each of the enriched pathway ontology terms across KEGG, Wikipathways and Reactome.

**Data File S10: Composition of cell types in each tissue.** Proportion of cells observed for each cell type and condition in each of the single cell datasets.

**Data File S11: Correlation between disease progression and healthy cell type program.**

## SUPPLEMENTARY MATERIALS

## SUPPLEMENTARY TABLES

**Supplementary Table 1.** Description of scRNA-seq datasets analyzed. We report the tissue of origin, number of cells, number of individuals and number of cell type programs analyzed for each single-cell dataset analyzed.

| **Tissue** | **# of cells** | **# of individuals** | **# of cell types** |
| --- | --- | --- | --- |
| **PBMC** (Travaglini et al) | 4,640 | 2 | 6 |
| **PBMC** (Zheng et al) | 68,551 | 8 | 6 |
| **Cord Blood** | 263,828 | 8 | 6 |
| **Bone Marrow** | 283,894 | 8 | 6 |
| **Brain** | 47,509 | 3 | 9 |
| **Kidney** | 40,268 | 13 | 24 |
| **Liver** | 13,340 | 4 | 12 |
| **Lung** | 31,644 | 10 | 19 |
| **Heart** | 287,269 | 7 | 12 |
| **Colon** | 110,373 | 12 | 20 |
| **Adipose** | 11,184 | 3 | 13 |
| **Skin** | 71,864 | 9 | 13 |
| **Colon (healthy + disease)** | 287,269 | 20 (healthy), 16 (disease) | 20 |
| **MS brain (healthy + disease)** | 48,918 | 9 (healthy), 12 (disease) | 12 |
| **Alzheimer's brain (healthy + disease)** | 70,634 | 24 (healthy), 24 (disease) | 8 |
| **Asthma lung (healthy + disease)** | 67,078 | 42 (healthy), 12 (disease) | 26 |
| **Idiopathic pulmonary fibrosis lung (healthy + disease)** | 114,396 | 10 (healthy), 20 (disease) | 19 |
| **COVID-19 BAL (healthy + disease)** | 43,930 | 3 (healthy), 6 (disease) | 10 |

**Supplementary Table 2.** Diseases and complex traits analyzed. We analyzed 60 diseases and complex traits with genetic correlation <= 0.9 and report the publication and sample size of each study.

| **Trait category** | **Trait** | **Source** | **Sample size (N)** |
| --- | --- | --- | --- |
| **Blood cell traits** | Lymphocyte percentage | UK Biobank | 444502 |
|  | Monocyte percentage | UK Biobank | 439938 |
|  | Platelet count | UK Biobank | 444382 |
|  | Red blood cell count | UK Biobank | 445174 |
|  | Red blood cell volume | UK Biobank | 442700 |
|  | Eosinophil count | UK Biobank | 439938 |
|  | Basophil count | UK Biobank | 439938 |
|  | Neutrophil count | UK Biobank | 439938 |
|  | Mean corpuscular volume | UK Biobank | 442122 |
| **Urine biomarkers** | Creatinine | UK Biobank | 434158 |
|  | Vitamin D | UK Biobank | 415700 |
|  | Bilirubin | UK Biobank | 429423 |
|  | Alkaline phosphatase | UK Biobank | 433862 |
|  | Aspartate amino transferase | UK Biobank | 430982 |
|  | Total protein | UK Biobank | 397652 |
| **Autoimmune diseases** | Inflammatory bowel disease | de Lange et al 2017 | 59957 |
|  | Crohn's disease | de Lange et al 2017 | 40266 |
|  | Ulcerative colitis | de Lange et al 2017 | 45975 |
|  | Eczema | UK Biobank | 458699 |
|  | Hypothyroidism | UK Biobank | 459324 |
|  | Rheumatoid Arthritis | Okada et al 2014 | 37681 |
|  | Primary biliary cirrhosis | Cordell et al. 2015 | 13239 |
|  | Lupus | Bentham et al. 2015 | 14267 |
|  | Type 1 diabetes | Bradfield et al. 2011 | 26890 |
|  | All autoimmune traits | UK Biobank | 459234 |
|  | Celiac disease | Dubois et al. 2010 | 15283 |
|  | Alzheimer's disease | Jansen et al. 2019 | 450988 |
|  | Multiple Sclerosis | Sawcer et al. 2011 | 27148 |
| **Neurological/ Psychiatric** | Number of children | UK Biobank | 456500 |
|  | Anorexia | Boraska et al 2014 | 32143 |
|  | ADHD | Demontis et al 2019 | 55374 |
|  | Autism | PGC cross disorder group | 10263 |
|  | Sleep duration | Dashti et al 2019 | 446118 |
|  | BMI | UK Biobank | 458417 |
|  | Major depressive disorder | Wray et al. 2018 | 173005 |
|  | Neuroticism | Nagel et al. 2018 | 449484 |
|  | Smoking status | UK Biobank | 457683 |
|  | Years of education | UK Biobank | 454813 |
|  | Intelligence | UK Biobank | 117131 |
|  | Morning person | UK Biobank | 410520 |
|  | Insomnia | Jansen et al. 2019 | 385506 |
|  | Schizophrenia | SCZ Working Group 2014 | 70100 |
|  | SCZ v. BD | Ruderfer et al 2018 | 38855 |
|  | Bipolar disorder | PGC bipolar group 2011 | 16731 |
|  | Reaction time | Davies et al 2018 | 300486 |
|  | Age of first birth | Barban et al. 2016 | 222037 |
| **Cardiac related traits** | Coronary artery disease | Schunkert et al 2011 | 77210 |
|  | ECG rate | UK Biobank | 53777 |
|  | Atrial Fibrillation | Nielsen et al. 2018 | 1030836 |
|  | Systolic blood pressure | UK Biobank | 422771 |
|  | Diastolic blood pressure | UK Biobank | 422771 |
| **Lung traits** | Childhood-Onset-Asthma | Ferreira et al. 2019 | 314633 |
|  | FEV1adjFEVC (lung capacity) | UK Biobank | 371949 |
|  | Idiopathic Pulmonary Fibrosis | Allen et al. 2020 | 11259 |
| **Other traits** | Height | Lango, Allen et al 2010 | 131547 |
|  | Breast Cancer | UK Biobank | 459324 |
|  | BMI-WHR | UK Biobank | 458417 |
|  | Type 2 Diabetes | Morris et al 2012 | 6078 |
|  | Basal metabolic rate | UK Biobank | 354825 |
|  | General risk tolerance | Karlsson Linner et al 2019 | 466571 |

## SUPPLEMENTARY FIGURES

**Supplementary Fig. 1.**
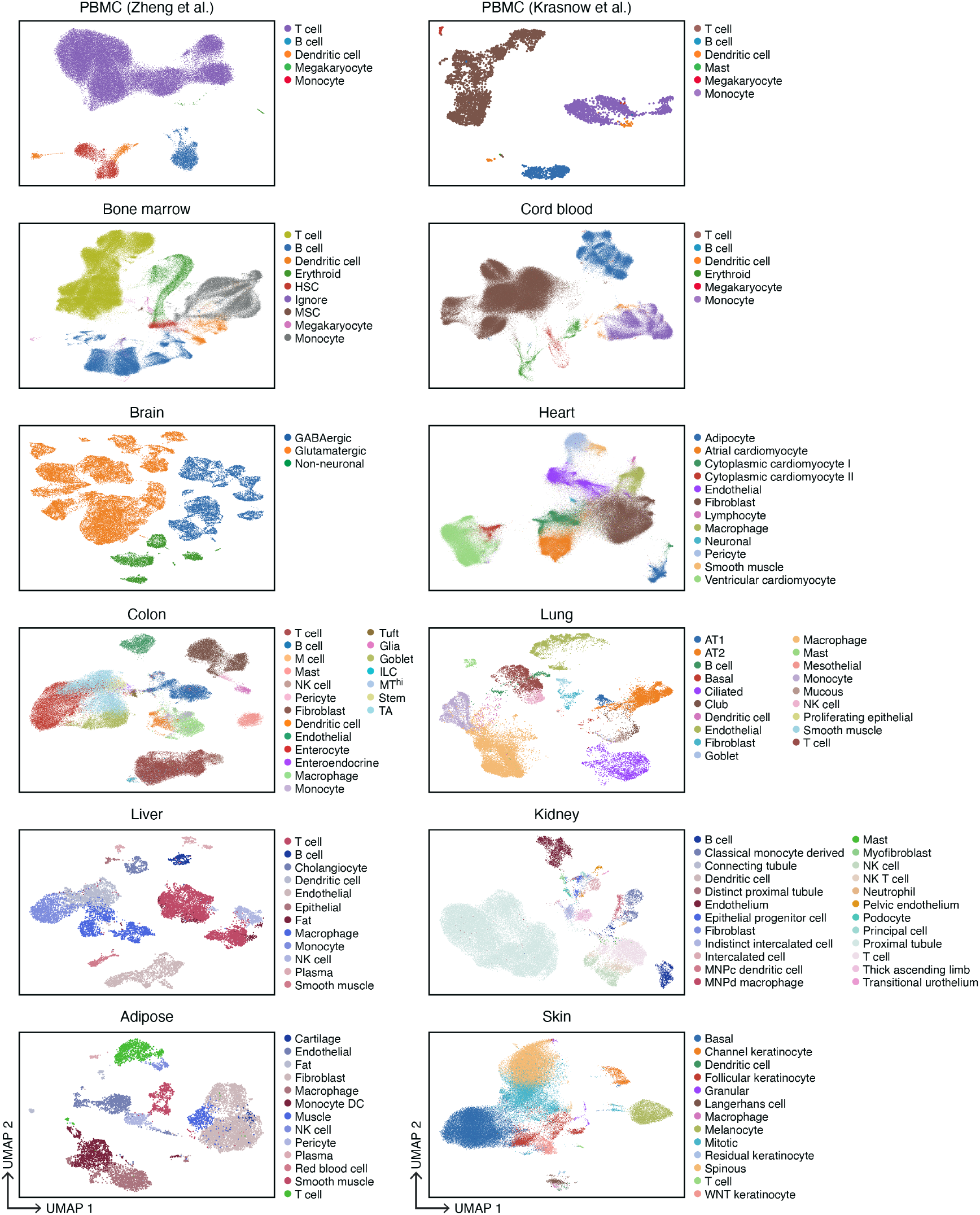
Single-cell RNA-seq datasets. UMAP embedding of scRNA-seq profiles (dots) colored by cell type annotations from 12 datasets (labels on top).

**Supplementary Fig. 2.**
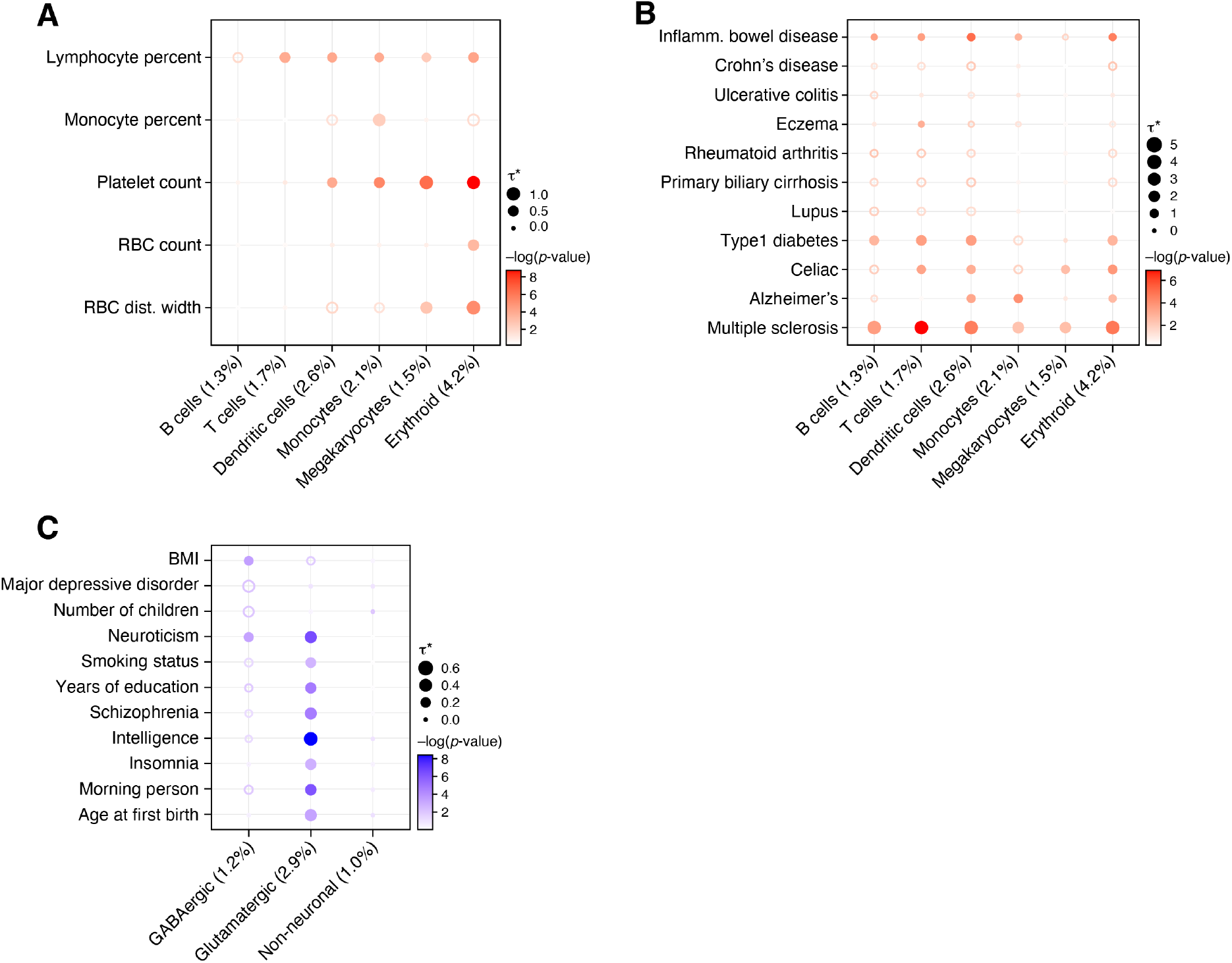
Standardized effect sizes of immune and brain cell type programs. Standardized effect size (!^∗^) (dot size) and significance (-log_10_(P-value), dot color) of the heritability enrichment of immune (**a,b**) or brain (**c**) cell type programs (columns) for blood cell traits (**a**), immune disease traits (**b**), or neurological/psychological related traits (**c**), based on SNP annotations generated with the Roadmap∪ABC-immune (**a,b**) or Roadmap∪ABC-brain (**c**) enhancer-gene linking strategy. Numerical results are reported in **data file S1**. Details for all traits analyzed are in **Supplementary Table 2.**

**Supplementary Fig. 3.**
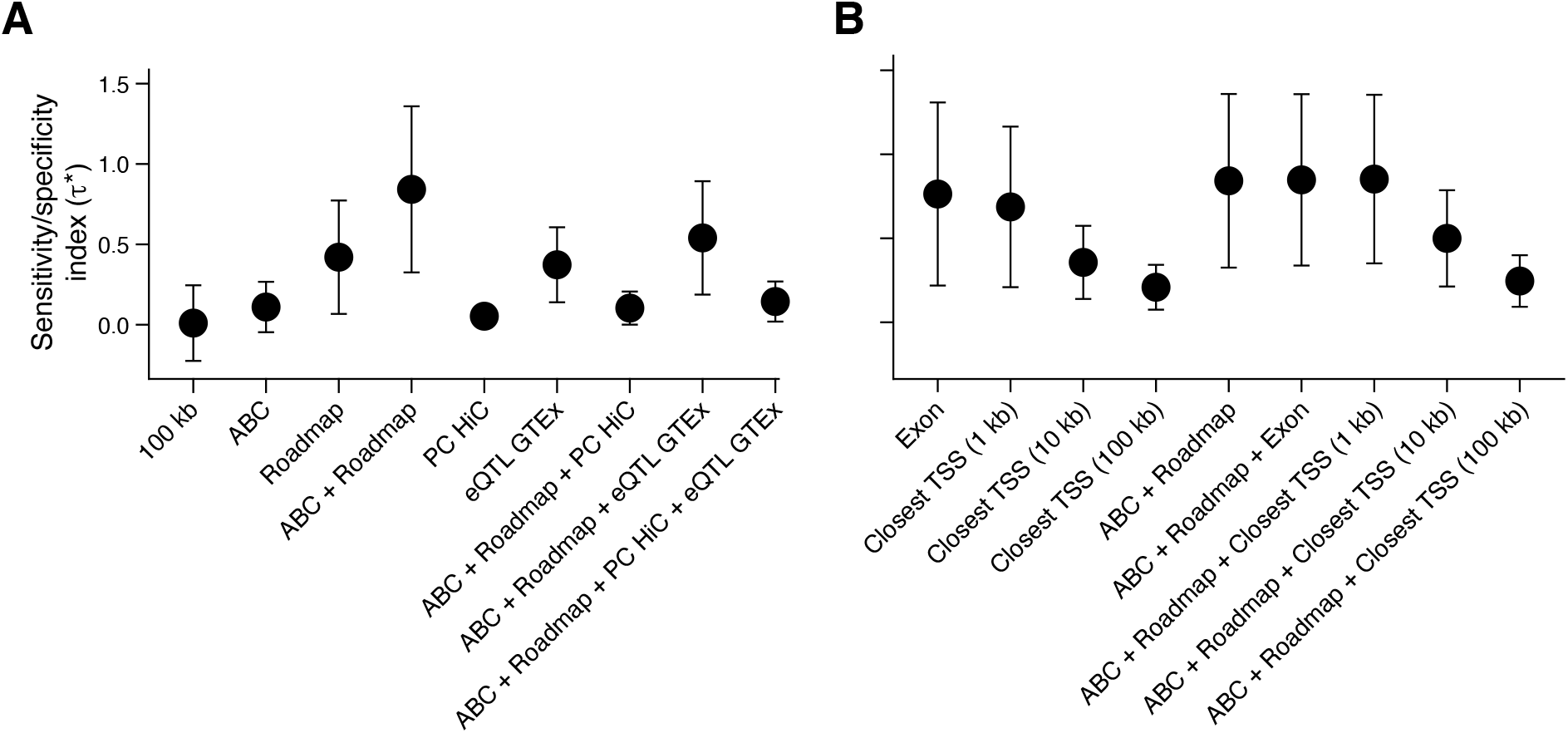
Roadmap∪ABC yields highest specificity of associations compared to other strategies. Specificity index (y axis, mean and s.e.) of immune programs and blood cell traits for different choices of regulatory regions linked to genes (*x* axis), including Roadmap∪ABC enhancer-gene strategy (ABC+Roadmap) and its constituent ABC and Roadmap strategies, promoter capture Hi-C (PC-HiC)^143, 144^ and eQTLs from the GTEx data^145^, and combination of Roadmap∪ABC with PCHiC (Roadmap+ABC+PCHiC), Roadmap∪ABC with eQTL (Roadmap+ABC+eQTLGTEx) and both PCHiC and eQTL (Roadmap+ABC+PCHiC+eQTLGTEx) (x axis, **a**), or closest TSS linking strategy between SNPs and genes at different distances (1kb, 10kb and 100kb), and their combinations with Roadmap∪ABC. Numerical results are reported in **data file S5**.

**Supplementary Fig. 4.**
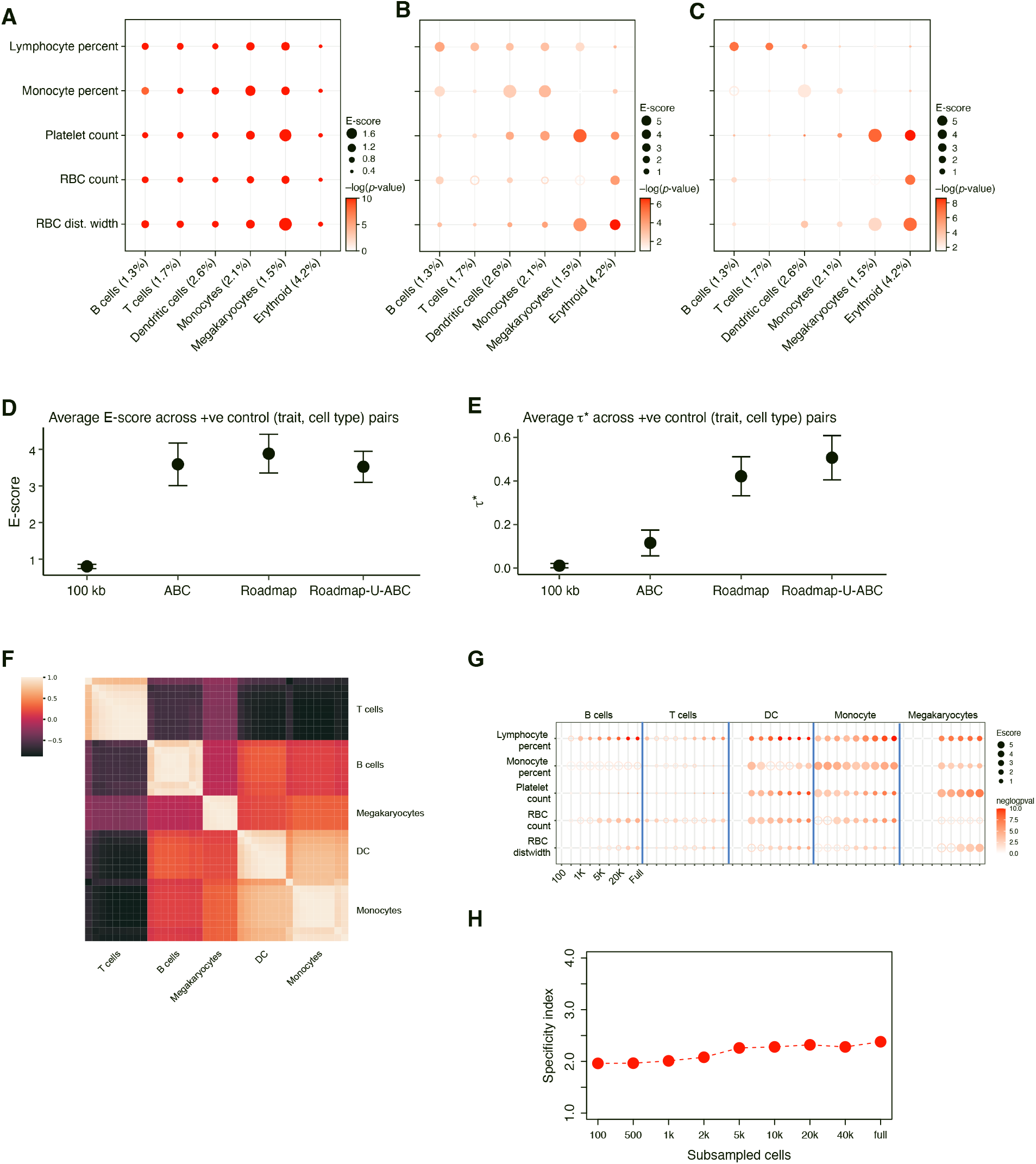
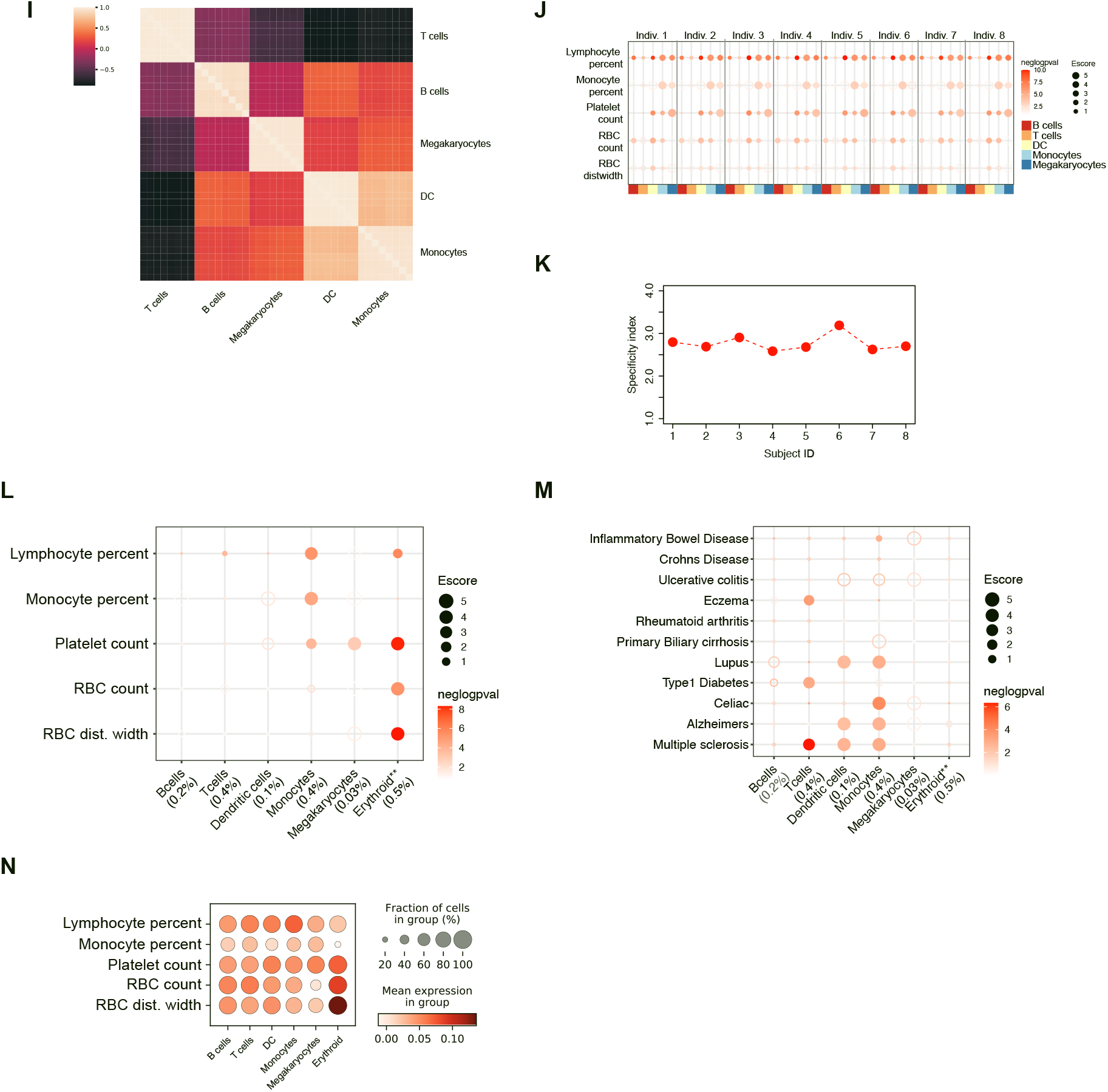
Benchmarking sc-linker across immune cell type programs and blood cell traits. **a-c**. Magnitude (E-score, dot size) and significance (-log_10_(P-value), dot color) of the heritability enrichment of immune cell type programs (columns) aggregated over 4 scRNA-seq datasets (PBMC (2), cord blood, and bone marrow) for 5 blood cell traits with SNP annotations combined with 100Kb (a), ABC-immune (b) or Roadmap-immune (c) strategies (compare to Roadmap∪ABC-immune strategy in **Fig. 2b**). **d,e.** Mean E-score (d) or average standardized effect size (!^∗^) (e) (y axis) for blood cell traits and immune cell type programs as in **Fig. 2b**, with SNP annotations combined with 100Kb, ABC-immune, Roadmap-immune or Roadmap∪ABC-immune strategy (x axis). Errors bars: 95% confidence intervals. **f.** Pairwise correlation heat map between all cell type programs computed for each sample separately. **g.** Magnitude (E-score, dot size) and significance (-log_10_(P-value), dot color) of the heritability enrichment of immune cell type programs constructed for each sample. **h.** Specificity index (y axis; see Methods) for immune cell type programs generated from each individual. **i.** Pairwise correlation heat map between all cell type programs computed for each dataset size separately. **j.** Magnitude (E-score, dot size) and significance (-log_10_(P-value), dot color) of the heritability enrichment of immune cell type programs constructed for each dataset size. **k.** Specificity index (y axis; see Methods) for immune cell type programs generated from subsampled PBMC scRNA-seq data at varying numbers of cells. **l,m.** Magnitude (E-score, dot size) and significance (-log_10_(P-value), dot color) of the heritability enrichment of immune cell type programs (columns) for 5 blood cell traits (l) and 11 autoimmune traits (m). **n.** Mean gene set expression score (dot color) from the baseline cell scoring approach. Comparison of panels l,m and n remains subjective, as the two metrics plotted (E-score/p.E-score in **l,m**; cell scores in **n**) are in different types of scoring schemes. Details for all traits analyzed are in **Supplementary Table 2.**

**Supplementary Fig. 5.**
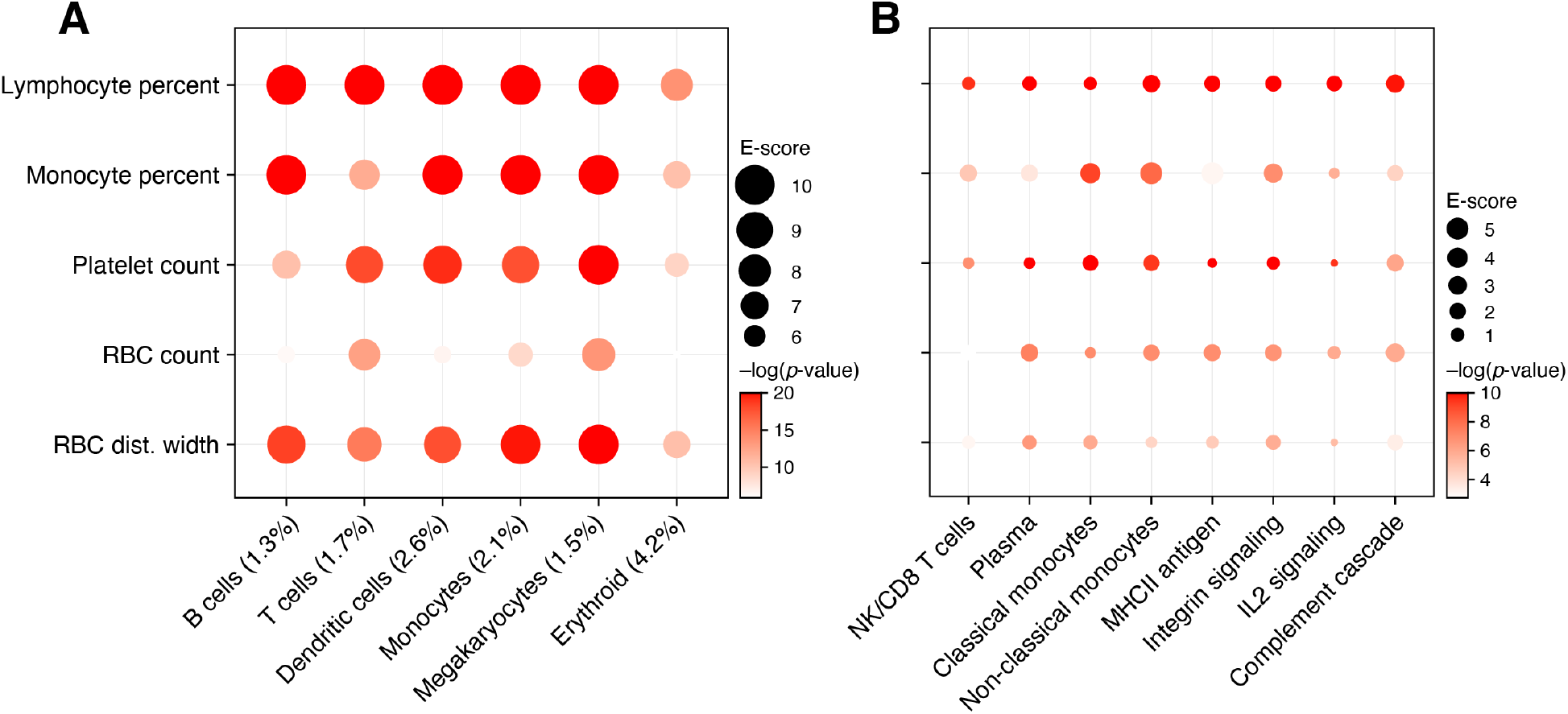
Analysis of functional enrichment of fine-mapped SNPs of immune cell type programs and heritability enrichment of immune cellular process programs. **a**. Functional enrichment of fine-mapped SNPs of immune cell type programs. Magnitude (Enrichment, dot size) and significance (-log_10_(P-value), dot color) of SNP annotations corresponding to immune cell type programs (using the Roadmap∪ABC-immune enhancer-gene linking strategy) with respect to functionally fine-mapped SNPs (from ref. ^146^). **b.** Heritability enrichment of cellular process programs for blood cell traits. Magnitude (E-score, dot size) and significance (-log_10_(P-value), dot color) of the heritability enrichment of immune cellular process programs (columns) and blood cell traits (rows). Details for all traits analyzed are in **Supplementary Table 2.**

**Supplementary Fig. 6:**
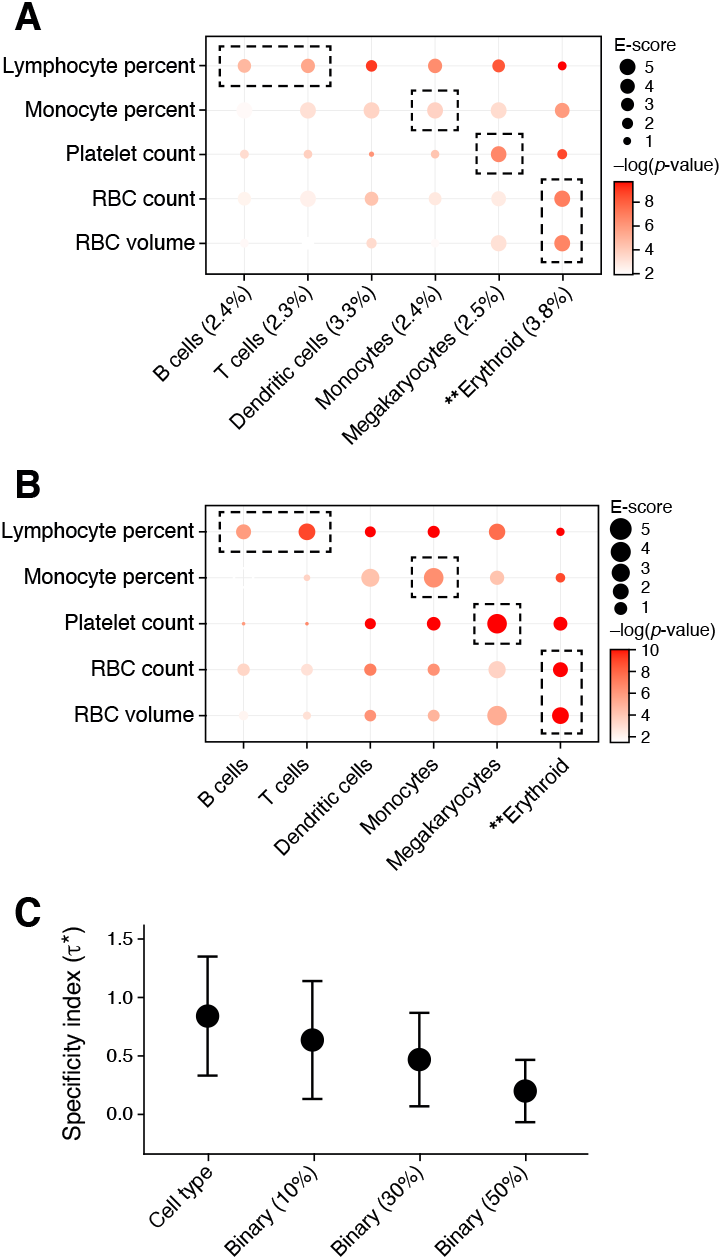
Evaluation of dichotomized gene programs. **a,b**. Enrichment in blood cell traits for binary and regular cell type programs. Enrichment (E score, dot size; and significance (-log_10_(P-value), dot color) for blood cell traits (rows) with cell type program defined by genes expressed in more than 10% of cells (**a**) or by our regular approach (**b**, as in **Fig. 2d**). The size of each corresponding SNP annotation (% of SNPs) is reported in parentheses. **c.** Regular cell type programs have a higher specificity than dichotomous ones. Specificity index metric (y axis, mean and s.e.) for blood biomarker and immune cell type programs defined by our regular approach (“cell type”) or by genes expressed in more than 10, 30 or 50% of cells of a given type (x axis). Numerical results are reported in **data file S6**.

**Supplementary Fig. 7.**
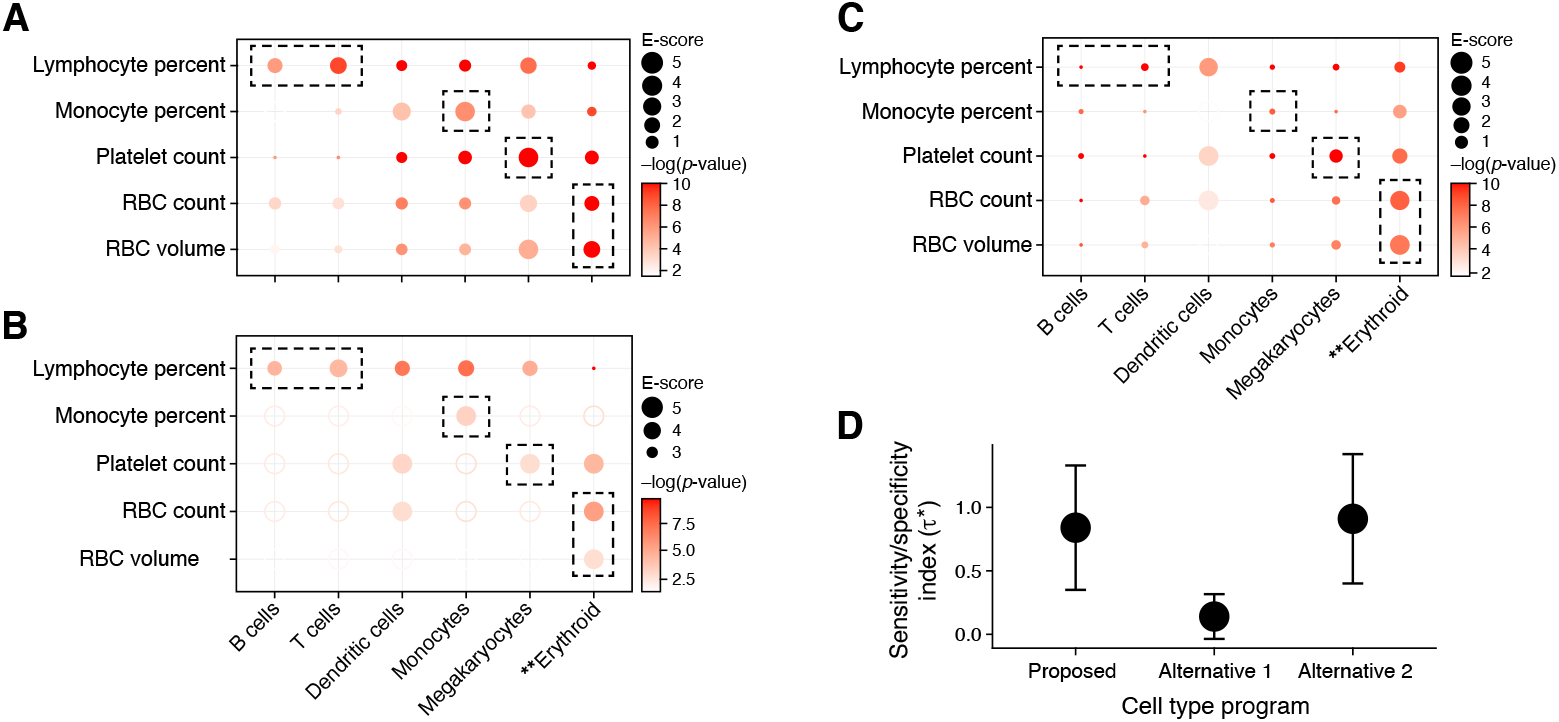
Evaluation of alternative approaches of gene program construction. **a-c**. Enrichment in blood cell traits for immune cell type programs defined in two different approaches. (**a**) Enrichment (E score, dot size; and significance (-log_10_(P-value), dot color) for blood cell traits (rows) with cell type programs (columns) defined either by genes differentially enriched in expression in a cell type compared to other genes in the same cell type (**a**), by genes differentially enriched in a cell type compared to their expression in other cell types (**b**, the primary analysis in this study), or by a combination of the previous two strategies (**c**). **d**. Sensitivity/specificity index of different approaches. Sensitivity/specificity index (y axis, mean and s.e.) for blood biomarker and immune cell type programs for the approaches in a-c. Numerical results are reported in **data file S6**.

**Supplementary Fig. 8.**
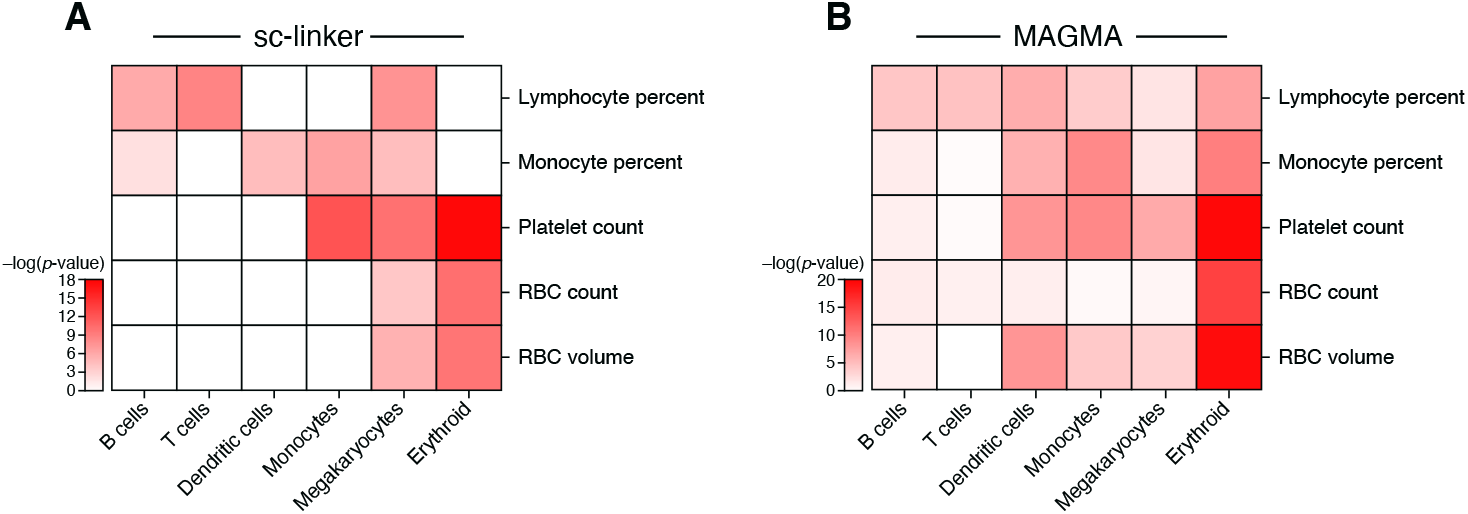
Comparison of sc-linker and MAGMA. Negative log p-value of immune cell type programs and blood cell traits for (**a**) E-score in sc-linker analysis, and (**b**) MAGMA gene-set level association analysis. Numerical results are reported in **data file S8**.

**Supplementary Fig. 9.**
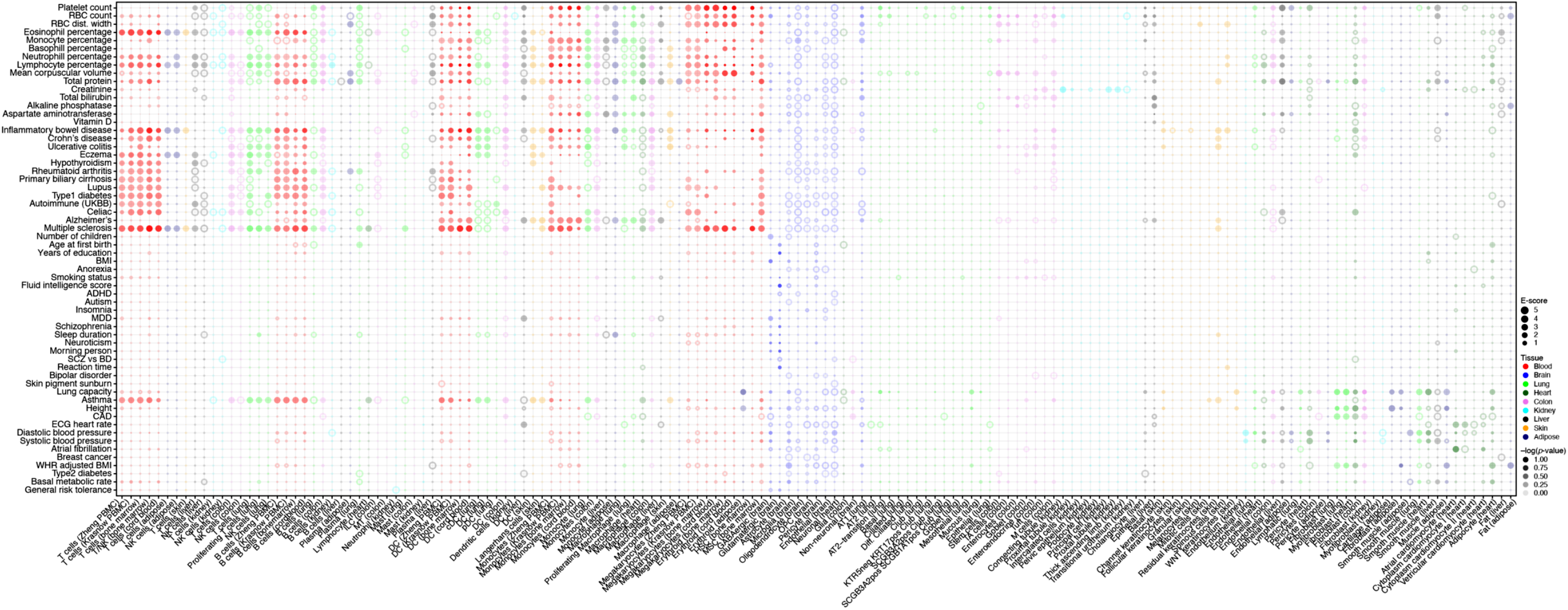
Linking cell type programs to diseases and traits across all analyzed tissues. Magnitude (E-score, dot size) and significance (-log_10_(P-value), dot color) of the heritability enrichment of cell type programs (columns) from each of nine tissues (color code, legend) for GWAS summary statistics of diverse traits and diseases (rows), based on the Roadmap∪ABC enhancer-gene linking strategy for the corresponding tissue. Details for all traits analyzed are in **Supplementary Table 2.** See **Data Availability** for higher resolution version of this figure.

**Supplementary Fig. 10.**
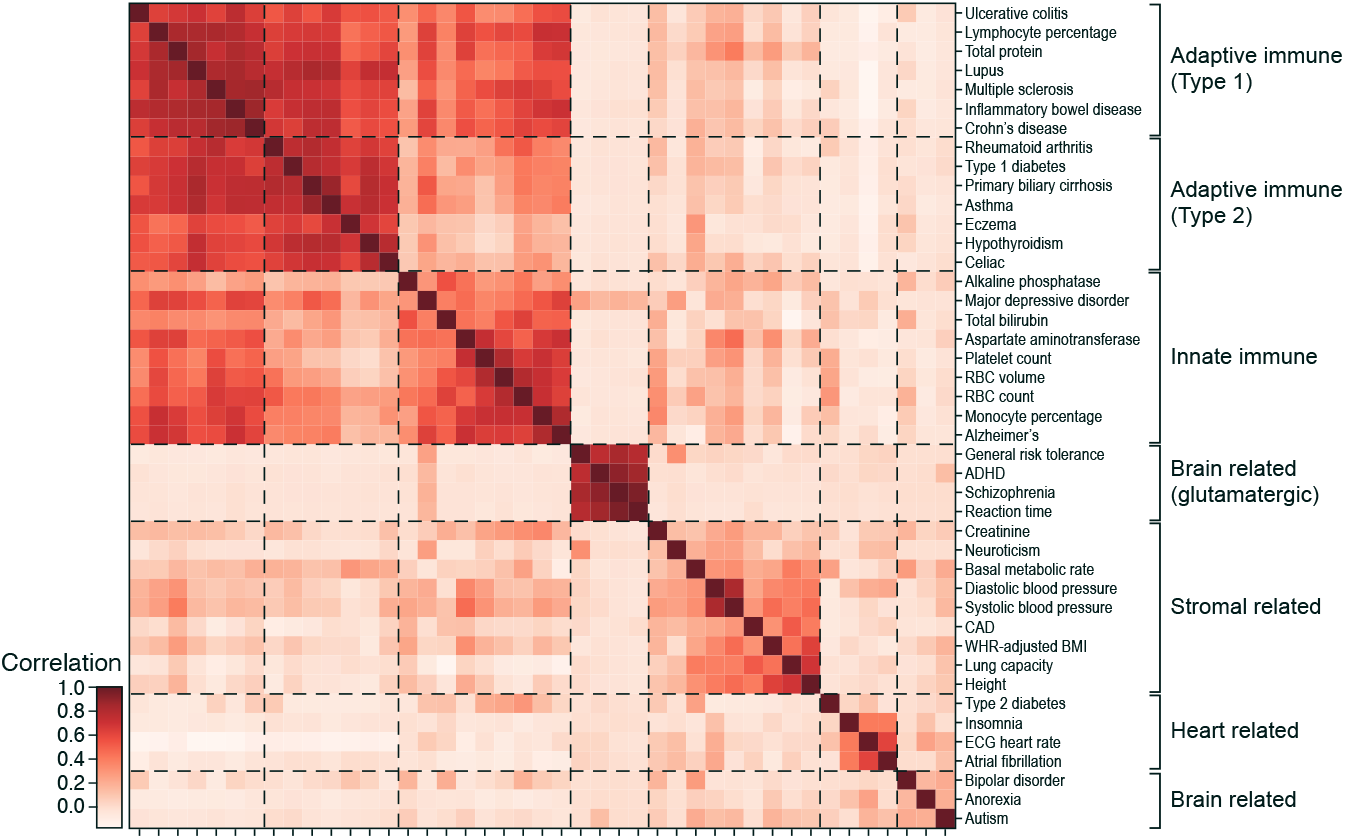
Cross trait analysis of cell type enrichments. Pearson correlation coefficient (colorbar) between the cell type enrichment profiles of each pair of traits (rows, columns), clustered (dashed lines) hierarchically. Trait clusters labeled by their overall cell type enrichments.

**Supplementary Fig. 11.**
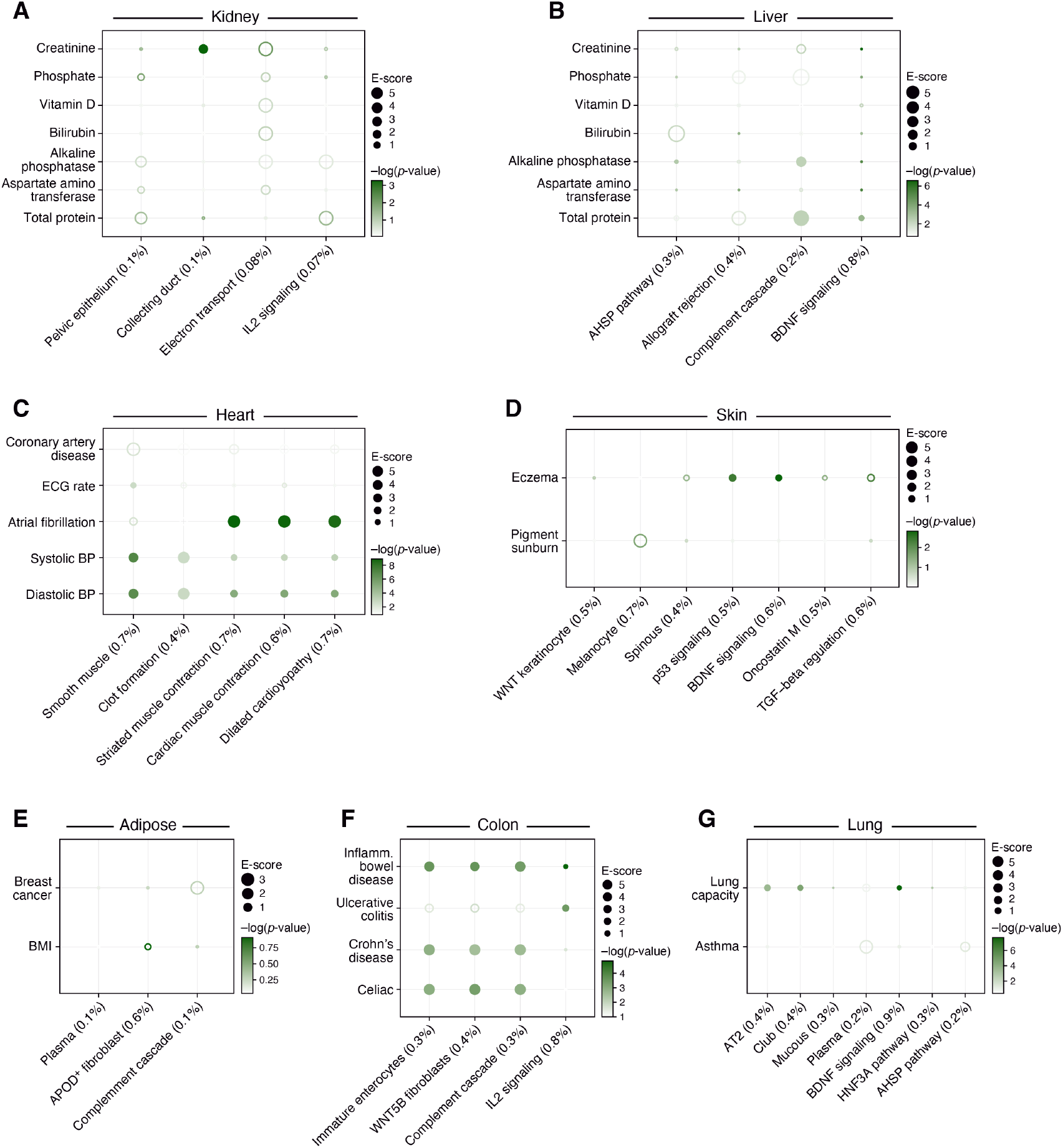
Linking cellular process programs to relevant diseases and traits in each of six tissues. Magnitude (E-score, dot size) and significance (-log_10_(P-value), dot color) of the heritability enrichment of cellular process programs (columns; obtained by NMF) in each of seven tissues (label on top) for traits relevant in that tissue (rows) using the Roadmap∪ABC strategy for the corresponding tissue. Details for all traits analyzed are in **Supplementary Table 2.**

**Supplementary Fig. 12.**
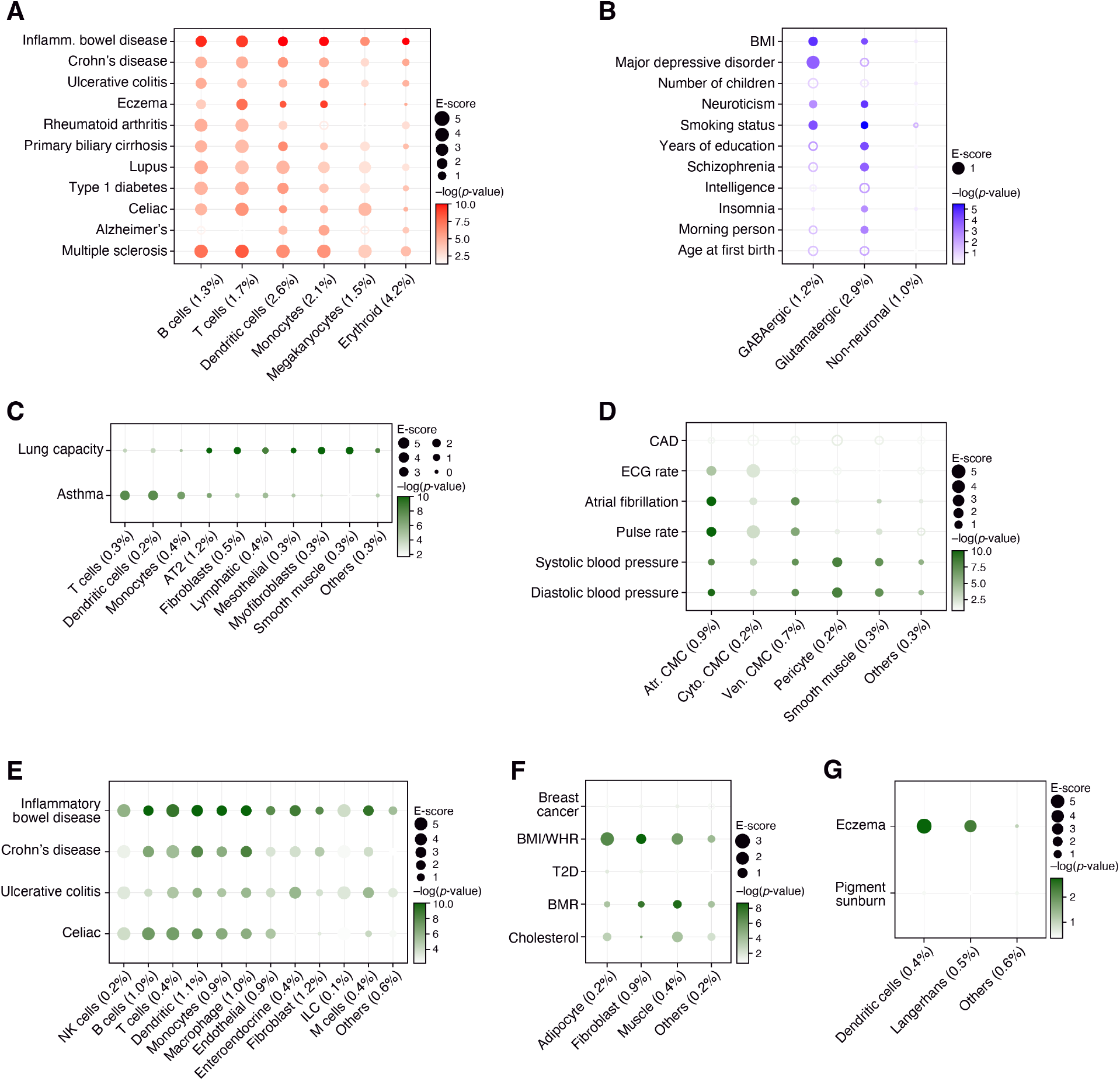
Analysis of cell type programs using a non-tissue-specific enhancer-gene linking strategy. Magnitude (E-score, dot size) and significance (-log_10_(P-value), dot color) of the heritability enrichment of immune (**a**), brain (**b**), lung (**c**), heart (**d**), colon (**e**), adipose (**f**) and skin (**g**) cell type programs (columns) for traits relevant in that tissue (rows) using a non-tissue-specific Roadmap∪ABC strategy. Details for all traits analyzed are in **Supplementary Table 2**.

**Supplementary Fig. 13.**
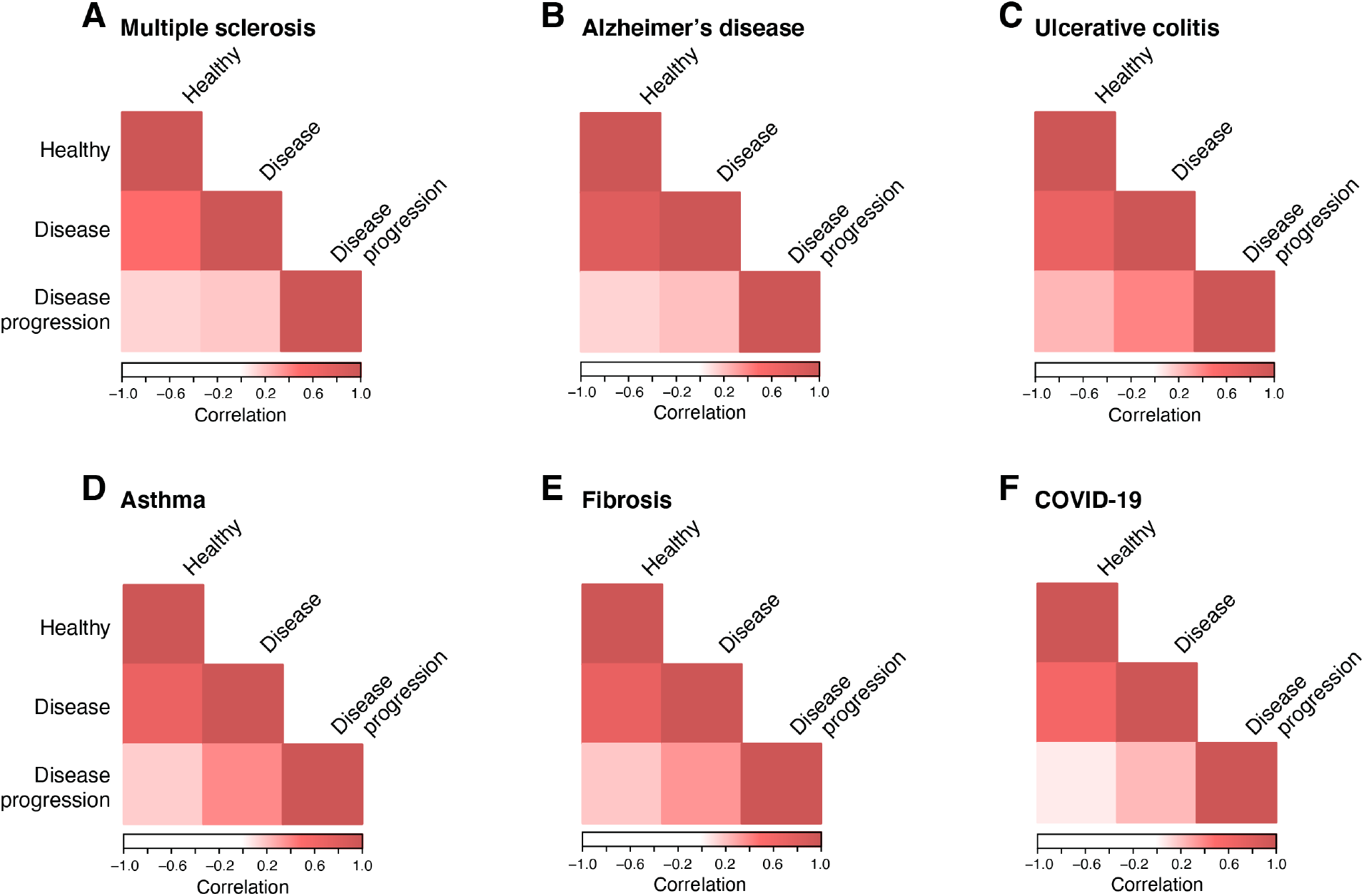
Disease progression programs have low correlations with healthy and disease cell type programs. Pearson correlation coefficient (color bar) of gene program membership vectors between healthy cell type, disease cell type and disease progression programs in scRNA-seq studies from a disease tissue (label on top) and the corresponding healthy tissue.

**Supplementary Fig. 14.**
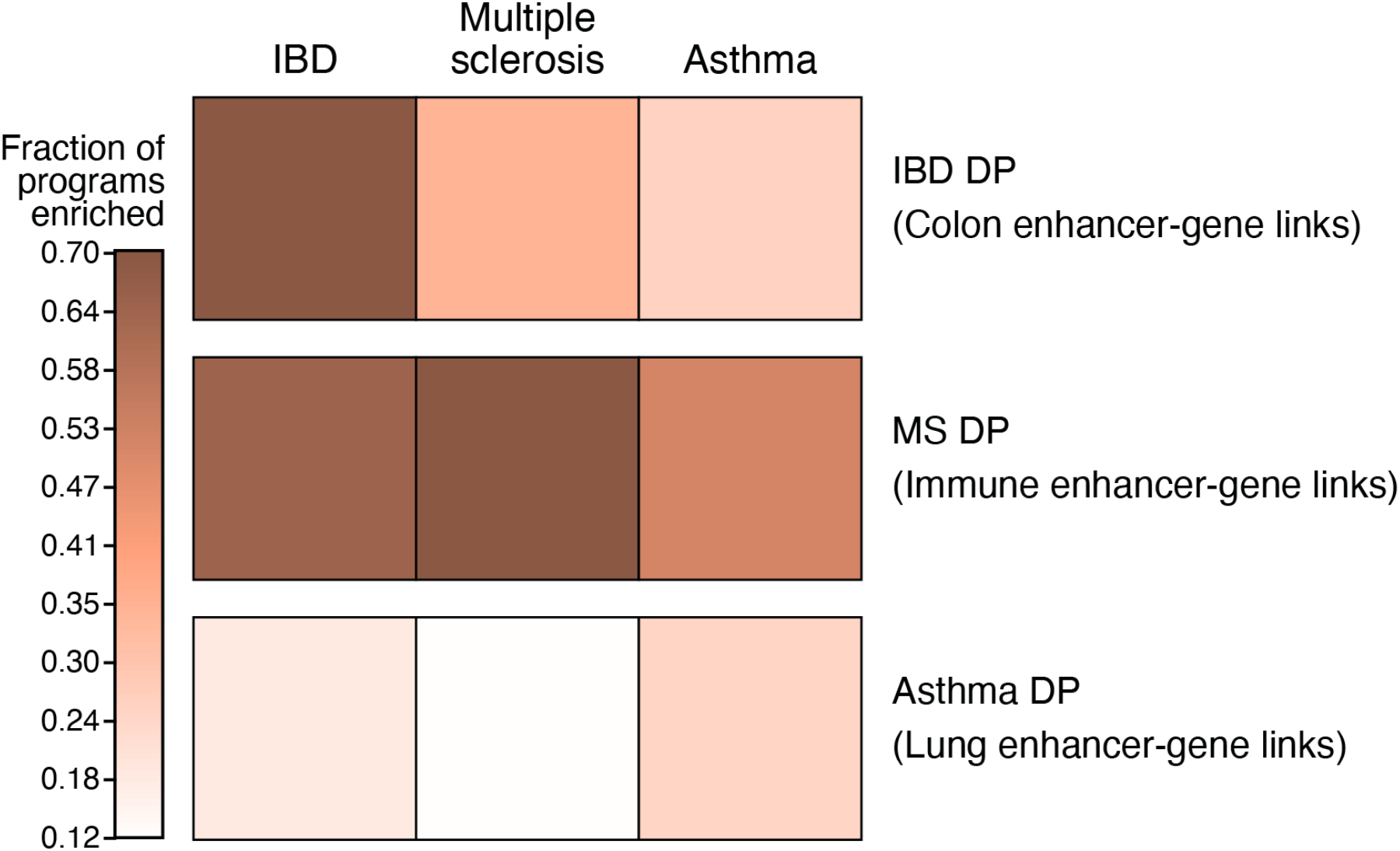
Disease specificity of disease progression programs. Proportion of disease progression programs with a -log_10_(P-value) of enrichment score (p.E-score) > 3 in IBD, MS and asthma GWAS summary statistics (column) for disease progression programs from IBD, MS and asthma (columns), when combined with tissue-specific Roadmap∪ABC (row).

**Supplementary Fig. 15.**
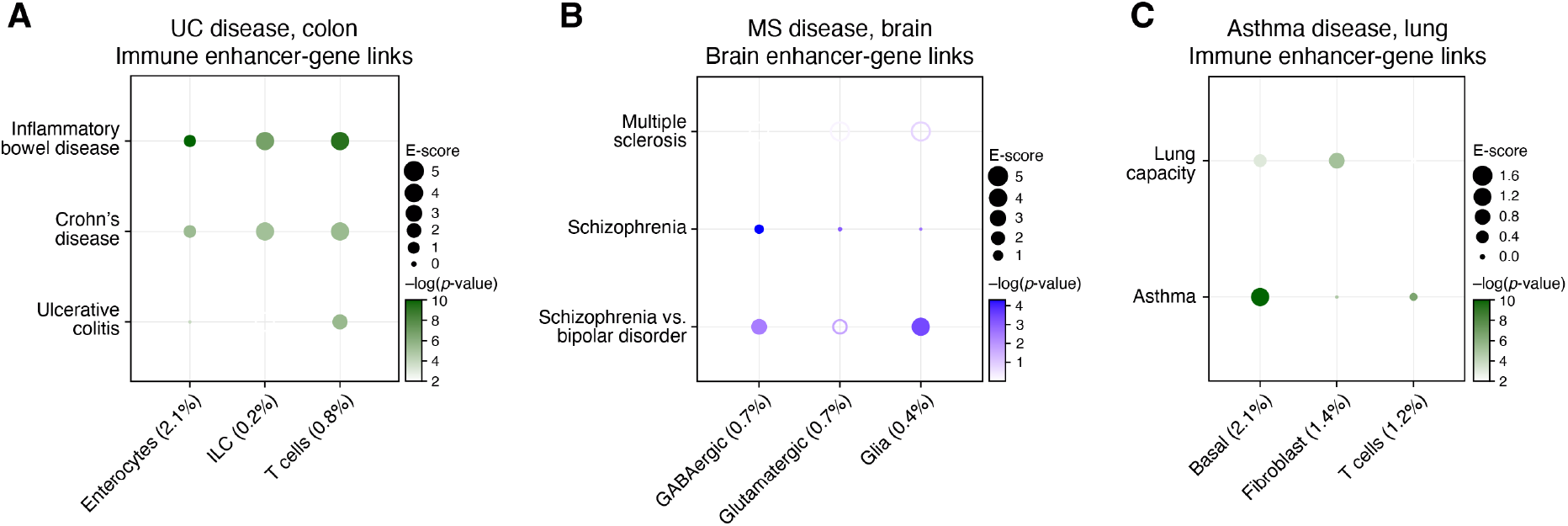
Analysis of disease progression programs using alternative Roadmap∪ABC enhancer-gene linking strategies. Magnitude (E-score, dot size) and significance (-log_10_(P-value), dot color) of the heritability enrichment of disease progression programs (columns) in UC (colon cells) using Roadmap∪ABC-immune (**a**), asthma (lung cells) using Roadmap∪ABC-immune (**b**), and MS (brain cells) using Roadmap∪ABC-brain (**c**). Details for all traits analyzed are in **Supplementary Table 2.**

**Supplementary Fig. 16.**
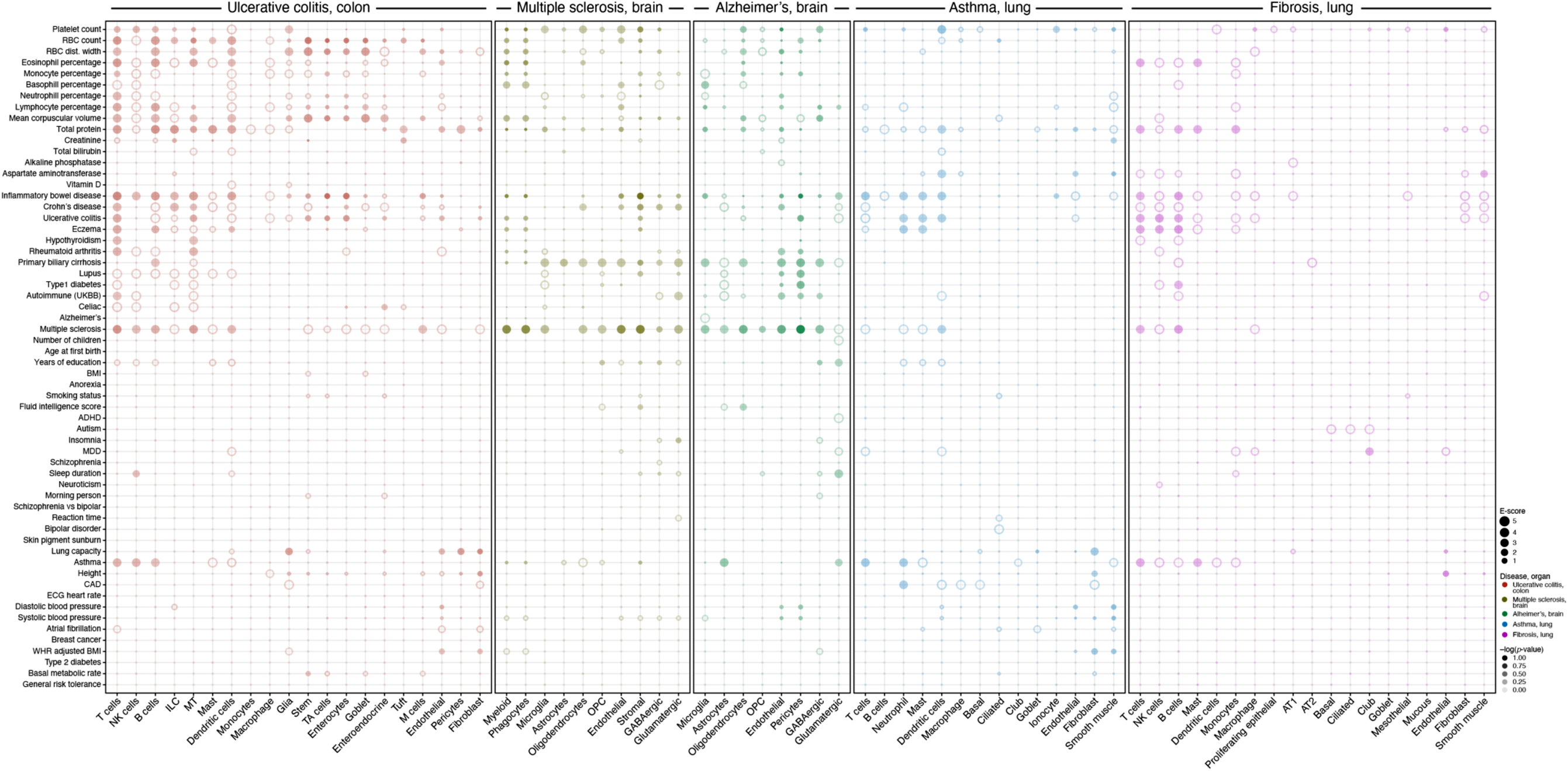
Analysis of disease progression programs across all tissues and traits. Magnitude (E-score, dot size) and significance (-log_10_(P-value), dot color) of the heritability enrichment of disease progression programs (columns) from UC, MS, Alzheimer’s, asthma and pulmonary fibrosis (labels on top, color code, legend), for GWAS summary statistics of diverse traits and diseases (rows), based on the Roadmap∪ABC enhancer-gene linking strategy for the corresponding tissue. Details for all traits analyzed are in **Supplementary Table 2.** See **Data Availability** for higher resolution version of this figure.

**Supplementary Fig. 17:**
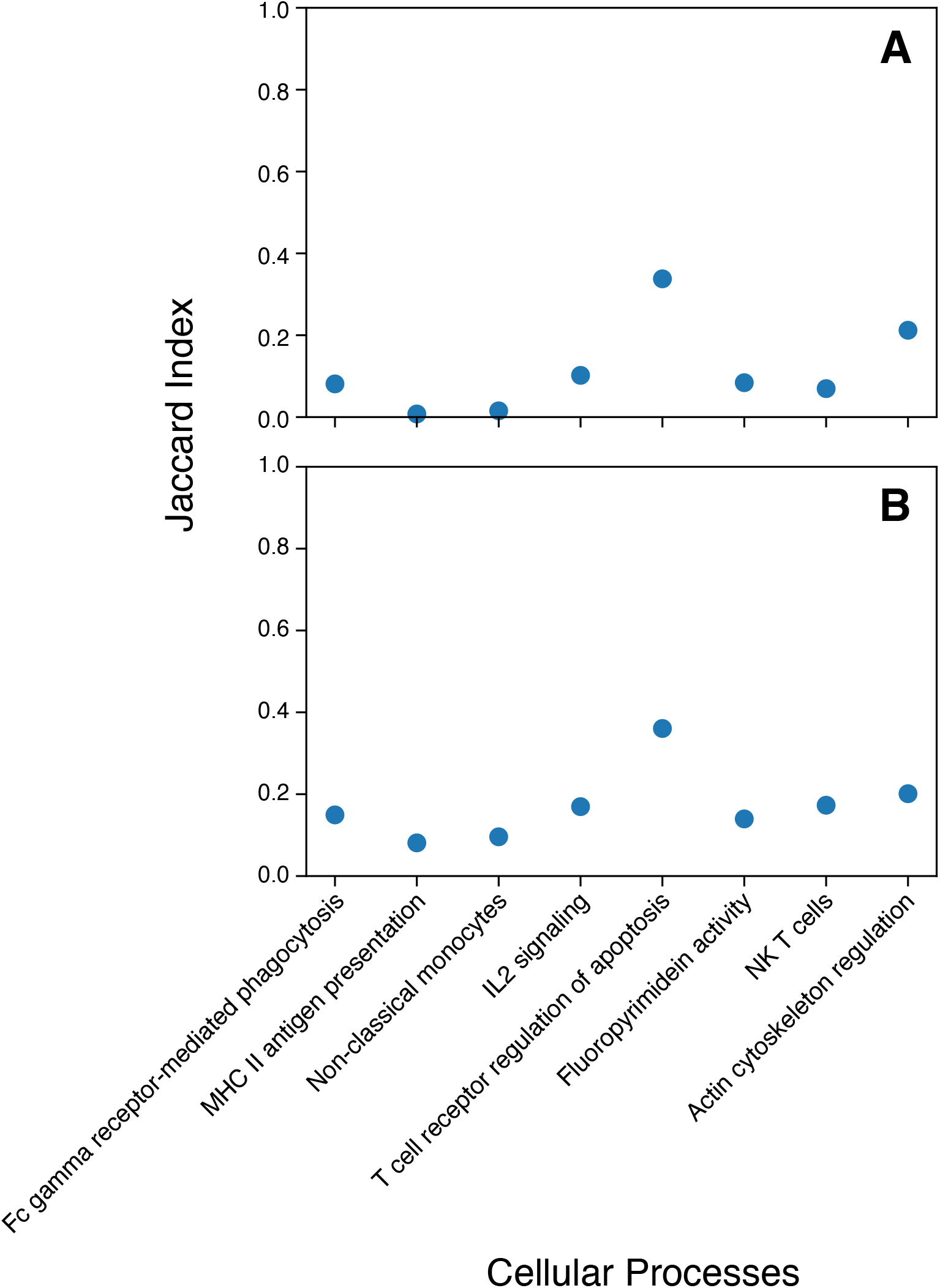
Top genes in blood cellular processes are neither highest expressed in cells nor in the tissue overall. Overlap (Jaccard index, y axis) between the top 200 genes in each blood cellular processes (x axis) and the highest expressed genes in the top 50 cells (based on the weight from the NMF decomposition) associated with the cellular process (**a**) or overall across the tissue (**b**).

### Supplementary Note

#### Extended analysis of disease critical brain cellular processes

The 12 brain cellular process programs showed that the significant enrichment of neuronal cell types above is primarily driven by finer programs reflecting neuron subtypes (**Fig. 3f, Table 1**). For example, the enrichment of GABAergic neurons for BMI was driven by programs reflecting LAMP5^+^ and VIP^+^ subsets; the respective top driving genes included *FLRT1* (for LAMP5^+^ neurons; ranked 1), whose absence reduces intercellular adhesion and promotes premature neuron migration^147^, and *TIMP2* (for VIP^+^ neurons; ranked 7), implicated in obesity through hypothalamic control of food intake and energy homeostasis in mice^148, 149^. Furthermore, the enrichment of GABAergic neurons for MDD reflects SST^+^ and PVALB^+^ subsets; the respective top driving genes included *PCLO* (for SST^+^ GABAergic neurons; ranked 2), and *ADARB1* (for PVALB^+^ neurons; ranked 4), encoding an RNA editing enzyme that can edit the transcript for the serotonin receptor 2C with a role in MDD^150^. We also observed enrichment in more specific cell subsets within the glutamatergic neurons (IT neurons were enriched for neuroticism, whereas L6 neurons were enriched for years of education and intelligence). Among inter cell type programs, electron transport cellular process programs (GABAergic and glutamatergic neurons) were enriched for several psychiatric/neurological traits, such as years of education, consistent with previous studies^77^, with the top driving genes including *ATP6V0B* and *NDUFAF3* (ranked 1, 4).

#### Role of healthy and disease progression T cells in Asthma

For example, healthy cell type and disease progression T cell programs were enriched in asthma, consistent with the contribution of T cell-driven inflammation to airway hyper-responsiveness and tissue remodeling^151^. From a pathway enrichment analysis, we identified that healthy T cell program overlapped with T cell receptor signaling, while the T cell disease progression program overlapped with RNA binding (**see data file S9**). These partially overlapping programs both included IL2 signaling pathway genes; IL2 is a T cell growth factor that increases airway response to allergens^152^ and drives differentiation of Th2 cells linked to asthma^153^.

#### Disease critical cell types in IPF and COVID-19

For IPF, a disease characterized by mucociliary dysfunction^154^, the mucous disease progression program was most enriched, and nominally significant (p = 0.04, not FDR significant), with top driving genes including *DSP* (ranked 1), a cell-cell adhesion molecule linked to tissue architecture in IPF lung^155^, and *MUC5B* (ranked 2), the well characterized genetic risk factor for IPF that likely increases mucinous expression in terminal airways of the lung^154^.

For severe COVID-19^156^, the macrophage disease progression program was enriched, and nominally significant (p = 0.01, not FDR significant), with top driving genes including key antiviral enzyme activators^157, 158^ *OAS3* and *OAS1* (ranked 1, 3), and *CCR5,* a chemokine receptor in which therapeutic intervention has been associated with improved prognosis in severe COVID-19 patients^159^. Further analyses of a meta-atlas of COVID-19 scRNA-seq in conjunction with COVID-19 GWAS data are described elsewhere^160^. Our nominally significant findings should be interpreted cautiously, but should become more powered as IPF and COVID-19 GWAS sample sizes grow.

